# Genome-wide association for loci influencing thermal performance curves in *Neurospora crassa*

**DOI:** 10.1101/2024.04.29.591604

**Authors:** Emmi Räsänen, Neda N. Moghadam, Karendeep Sidhu, Pauliina A. M. Summanen, Henna-Riikka Littunen, Tarmo Ketola, Ilkka Kronholm

**Author notes:** Corresponding author: Ilkka Kronholm Department of Biological and Environmental Science, University of Jyväskylä, P.O. Box 35, FI-40014 Jyväskylä, Finland.

## Abstract

Temperature poses a unique challenge to ectothermic species, as it affects all biochemical reactions in the cell and causes physiological stress. The effect of temperature on an organism can be described by a thermal performance curve (TPC), which displays organismal performance, such as growth rate, as a function of temperature. Previous studies on thermal performance have revealed different amounts of genetic variation and trade-offs in TPC shape and position within species and populations. However, very little is known about the genetic architecture of TPCs on the level of individual loci and alleles. We asked what is the identity of loci contributing to genetic variation in TPCs, and do the alleles exhibit trade-offs or thermodynamic scaling across the temperature range? We used genome-wide association mapping to find loci influencing growth rate at different temperatures and TPC traits in the filamentous fungus *Neurospora crassa*. We also evaluated the directions and magnitudes of allelic effects to investigate possible trade-offs. We observed both unique associations at specific temperatures, as some loci affected growth rate only at low, intermediate, or high temperatures, and associations that were shared across multiple temperatures. However, only weak evidence of trade-offs was detected, indicating that the evolution of TPCs in *N. crassa* is not constrained by allelic effects in opposite directions at hot and cold temperatures. Our findings indicate that trade-offs contribute little to variation in TPCs.

## Introduction

Among all of the abiotic stress factors, temperature poses a unique challenge for ectothermic organisms as temperature affects the stability of proteins and all biochemical reactions in the cell (Schulte, 2015; Arcus *et al*., 2016). Therefore evolutionary adaptation to higher temperatures may be more difficult than to other abiotic stresses. This is especially concerning today as global temperatures are increasing as a result of greenhouse gas emissions (IPCC, 2013). As some organisms may have no possibility to migrate into colder areas, they have to adapt to higher temperatures, or otherwise they may perish (Deutsch *et al*., 2008; Dillon *et al*., 2010; Araújo *et al*., 2013; Merilä and Hendry, 2014).

The response to temperature an ectothermic organism exhibits is called a thermal performance curve (TPC) (Huey and Kingsolver, 1989, 1993). This curve describes an aspect of organism’s performance, such as growth or fitness, as a function of temperature. Often TPCs are skewed towards colder temperatures, and they have an optimum temperature, after which performance rapidly falls at higher temperatures (Sinclair *et al*., 2016). As different species have TPCs with different optima and shapes (Maclean *et al*., 2019; Kontopoulos *et al*., 2020), genetics of the organism must be important for determining the TPC shape. Generally, most studies have detected genetic variation in thermal performance within species, although the extent of this variation differs between species and populations (Krenek *et al*., 2011; Logan *et al*., 2018, 2020; Martins *et al*., 2019; Moghadam *et al*., 2020; Querns *et al*., 2022). Genetic variation is mostly restricted to certain aspects of temperature performance, for example, many studies report that majority of genetic variation is in TPC elevation rather than in the shifting optimum temperature (Shama *et al*., 2011; Klepsatel *et al*., 2013; Latimer *et al*., 2015; Bartheld *et al*., 2017; Moghadam *et al*., 2020). However, a meta-analysis of selection experiments has revealed evidence of trade-offs when a population adapts to a new temperature optimum (Malusare *et al*., 2023).

Since the activation energy of every biochemical reaction in the cell scales with temperature, thermodynamic effects must also influence the characteristics of TPCs (Angilletta *et al*., 2010; Asbury and Angilletta Jr., 2010). Thermodynamic constraints and trade-offs are expected to affect the evolution of species’ thermal performance since the enzymatic reactions slow down at colder temperatures (Kontopoulos *et al*., 2020). Conversely, the rate of biochemical reactions and also the organism’s performance should be higher at warmer temperatures. Based on this specialisation and thermodynamics, cold-adapted species should have lower fitness at warmer temperatures and vice versa, which is observed as a optimum-shift between colder and warmer temperatures in TPCs (Malusare *et al*., 2023). Cold-adapted species could also be constrained in their ability to enhance performance to the same level as warm-adapted species through evolution, unless there is a shift towards higher temperatures in TPC, also know as the "hotter is better" hypothesis (Angilletta *et al*., 2010). The evidence for thermodynamic constraints is mixed, as some studies have reported larger effects (Frazier *et al*., 2006; Sørensen *et al*., 2018), and others only minor effects (Kontopoulos *et al*., 2020; Malusare *et al*., 2023).

While the genetics of TPCs has been investigated with quantitative genetic methods and QTL studies that have a low mapping resolution (Latimer *et al*., 2015), we understand the effects of individual genes on TPCs poorly. What is the identity of genes that influence TPCs? How do allelic effects scale with temperature? For example, are there loci for which allelic effects scale thermodynamically, and other loci that affect growth only at specific temperatures? Moreover, the theory of TPC evolution often assumes that there are trade-offs between performance at higher and lower temperatures (Angilletta Jr., 2009; Angilletta *et al*., 2010), but can this be observed on the level of genes or alleles?

In order to better understand the genetic architecture of TPCs, we performed a genome-wide association study for growth rate at different temperatures in the filamentous fungus *Neurospora crassa*. This ascomycete fungus is a genetic model system with both asexual and sexual life cycle (Roche *et al*., 2014). It can be propagated clonally through asexual spores, or mating can be induced between individuals of different mating types. In nature, *N. crassa* is a subtropical species, that decomposes dead plant material. We used strains collected from Southeastern United States and the Caribbean (Ellison *et al*., 2011; Palma-Guerrero *et al*., 2013), and made crosses among some of the strains to generate a nested association mapping population. We have previously investigated the genetic variation in TPCs for this population on the phenotypic level (Moghadam *et al*., 2020), and here we use genotypic information for these lines to map loci that influence growth at different temperatures.

## Materials and methods

### Neurospora crassa strains

For nested association mapping, we used the strains described in Moghadam *et al*. (2020). Briefly, the strains used in this study included natural strains collected from Lousiana (USA), Caribbean, and Central America (Ellison *et al*., 2011; Palma-Guerrero *et al*., 2013). Sampling locations are shown in figure S1. We have made crosses between some of the natural strains to create additional strains for nested association mapping (Table 1). Strains belonging to a particular family are F_2_ offspring of the parents. In total there were 434 strains in this study. Full list of the strains is presented in supplementary table S1.

**Table 1:** Strains for nested association mapping panel used in this study. FGSC = Fungal Genetics Stock Center.

| Population | *n* | Parents | Source |
| --- | --- | --- | --- |
| Natural strains | 118 | – | FGSC |
| Family A | 94 | 10948 × 10886 | (Moghadam *et al.*, 2020) |
| Family B | 50 | 10932 × 1165 | (Moghadam *et al.*, 2020) |
| Family C | 50 | 4498 × 8816 | (Moghadam *et al.*, 2020) |
| Family D | 52 | 3223 × 8845 | (Moghadam *et al.*, 2020) |
| Family G | 70 | 10904 × 851 | (Moghadam *et al.*, 2020) |
| Total | 434 | – | – |

#### Genotyping

A dense genotyping is needed for association mapping, but high density SNP chips are not available for *N. crassa*, and sequencing the genomes of all the 316 offspring from the families generated by crosses was not possible. Therefore, we employed a strategy where we used the full genome sequencing data only for the 118 natural strains. Then we used RAD-sequencing (Baird *et al*., 2008) to obtain a moderate number of SNP genotypes for each offspring. We could then use the RAD markers to find the locations of recombination breakpoints. Because we know the full genome sequences of the parents, we could then infer the genome sequences for the offspring based on their RAD markers.

##### Genome re-sequencing of strains from natural populations

Some of the strains from natural populations had already been sequenced in previous studies, 23 strains had been sequenced by Zhao *et al*. (2015), 33 strains by Villalba de la Peña *et al*. (2023), and 61 strains were sequenced in this study. Including the reference genome of strain 2489, this comes to 118 re-sequenced strains. See table S2 for list of the sequenced strains and alignment metrics.

We used the same methods for DNA extraction, sequencing, and SNP calling as in Villalba de la Peña *et al*. (2023). Briefly, DNA was sent to Novogene (Cambridge, UK) for sequencing, using Illumina 150 bp paired-end reads. Reads were mapped against the reference genome (version 12) using BWA-MEM (Li, 2013). SNPs were called using Haplotypecaller in GATK (McKenna *et al*., 2010), and filtered using Wormtable (Kelleher *et al*., 2013) and custom python scripts. For a SNP to be included it had to have read depth *≥* 5 in the focal sample, and mean *≥* 5 across all samples. The same procedure was used with genotype quality *≥* 30, and mapping quality *≥* 40. We performed genotyping in diploid mode, even if *N. crassa* is haploid, and excluded heterozygous sites. Based on our experience, sites appearing as heterozygous in a haploid organism represent read mapping problems, see also Kronholm *et al*. (2017). If a site did not pass the filters for a given sample, the genotype of that sample was assigned as missing data. See methods in Villalba de la Peña *et al*. (2023) for details.

##### RAD-sequencing

We genotyped the 315 offspring of the crosses using RAD-sequencing (Baird *et al*., 2008). We extracted DNA from the strains with phenol-chloroform extraction and sent the samples for RAD-sequencing by Floragenex Inc. (Oregon, USA), using a single digest with Pst I, barcode multiplexing, and Illumina sequencing with 100 bp read length.

The RAD-reads were cleaned and demultiplexed with the STACKS program (Catchen *et al*., 2011). Across all RAD-samples 94.6% of the reads were retained after filtering reads of low-quality, or where the barcode could not be identified. Reads were aligned to the *Neurospora crassa* reference genome using BWA-MEM (Li, 2013), and the resulting bam files were sorted with samtools. We added read groups and performed indel realignment with GATK (McKenna *et al*., 2010) with parents and offspring of each family together, and then used Unified Genotyper in GATK for genotype calling with parameters: heterozygosity = 0.001, ploidy = 2, and minimum base quality score = 10. The resulting vcf files were processed using custom R scripts. For each family, we filtered out loci that had *<* 20 individuals genotyped, loci that had genotype quality *<* 30, mean read depth *≤* 10 across the family, minor allele frequency *≤* 0.2, or had an unbalanced allelic depth ratio (log of ratio *< −*1 or *>* 1). Further criteria included that at least one of the parents was genotyped and parental genotypes were different, neither parent was called heterozygous, and proportion of individuals called as heterozygous was *<* 0.1. For the remaining loci, all genotype calls with read depth *≤* 5 were set to missing data. There were also some loci that had been called as both a SNP and an indel, at the same position. In these cases only the indels were kept, as these polymorphisms likely represented complex variants. After quality filtering there were 14 445 loci left for family A, 13 523 loci left for family B, 17 297 loci left for family C, 16 314 loci left for family D, and 11 261 loci left for family G.

Criteria for filtering RAD-loci was assessed by constructing genetic maps for each family using the R package ASMap (Taylor and Butler, 2017). We used the kosambi mapping function, and a p-value cut off of 10*^−^*^6^ for evidence of linkage. We observed that leaving markers with bad quality in the dataset produced genetic maps with excessive numbers of double cross-overs, or large local segregation distortion resulting from genotyping errors. For the filtered dataset some double cross-overs were still observed (Figure S2), while it is possible that some were due to genotyping error, most are likely real events. Marker order was generally concordant with the reference genome (Figure S3).

##### Inference of full SNP genotypes for the offspring

Eleven to seventeen thousand markers obtained with RAD-sequencing is not enough for high resolution association mapping, but they allowed us to identify the locations of recombination events in the offspring genomes. We used locations of recombination breakpoints to infer the genotypes of the offspring for all of the SNPs genotyped for the natural strains. We classified the genome of each offspring into segments inherited from the two different parents (e.g. Figure S4), SNPs for chromosome ends were classified according to the first or the last RAD-marker. Where recombination breakpoints occurred, those SNPs falling between the two RAD-markers flanking the breakpoint were set as missing data. Because some individuals had a lot of missing data for the RAD-markers, we were conservative and segments with missing RAD-data were inferred as missing for the SNP genotypes. The median distance between two RAD markers flanking the recombination break-points was 11.5 kb. Because *N. crassa* is haploid and the parent genotypes are known, there is no ambiguity in inferring the genotypes of the offspring due to heterozygosity or phasing issues. After inferring offspring SNP genotypes, we created the final genotype dataset for association mapping by filtering out SNPs that were not biallelic, had minor allele frequency *<* 0.01, or were not mapping to the seven *N. crassa* chromosomes. This resulted in 1 473 869 SNPs for the nested association mapping population of 434 strains.

#### Phenotyping

The association mapping panel strains have been previously phenotyped for growth rate at six different temperatures: 20, 25, 30, 35, 37.5, and 40 °C (Moghadam *et al*., 2020). Growth rate was measured as the expansion of the mycelium (mm*/*h) in a linear tube. We used asexual spores to propagate the strains, so the same individuals could be measured multiple times and in different environments. Mean of three replicates was used as the phenotypic value for a given strain in a given temperature. See Moghadam *et al*. (2020) and Kronholm *et al*. (2016) for details about growth rate measurements.

#### Statistical analysis

All statistical analyses were performed with the R environment version 3.6.0 (R Core Team, 2019) and the R package ’ggplot2’ (Wickham, 2016) was used for plotting. To obtain estimate of the TPC for each strain we fitted a natural spline through the data of each strain, and extracted optimum temperature and maximum growth rate from these fits, see Moghadam *et al*. (2020) for details.

##### Linkage disequilibrium

To evaluate linkage disequilibrium (LD) between markers in our mapping population, we used only the 118 strains that belong to the natural populations. To evaluate how LD decays over a long distance, we first sampled 10 000 pairs of SNPs randomly from each chromosome for which we calculated LD, for a total of 70 000 pairwise comparisons. To evaluate LD on finer scales we sampled 10 sets of SNPs spaced evenly across each chromosome. For each set we took the 99 consecutive SNPs from the focal SNP and calculated LD for all unique pairwise comparisons for these 100 SNPs. This resulted in 4950 comparisons for each SNP set, and for 10 sets per chromosome and seven chromosomes this comes to 346 500 comparisons in total.

Coefficient of LD was calculated following Slatkin (2008) as *D* = *p_AB_− p_A_p_B_*, where *p_AB_* is the haplotype frequency of haplotype AB and *p_A_*and *p_B_*are allele frequencies for *A* and *B* respectively. Then LD was calculated as

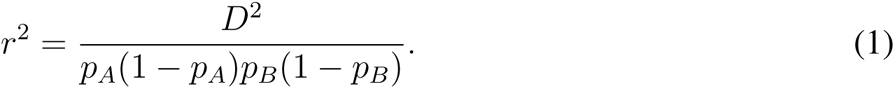

##### Genome-wide association

For genome-wide association we used a mixed model association implemented in the GAPIT 3 program (Wang and Zhang, 2021). The model accounts for population structure and relatedness among individuals (Zhang *et al*., 2010). The basic model was

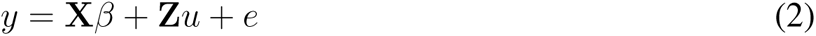

where *y* is a vector of phenotypic observations, *β* is a vector of fixed effects including effects of the genetic marker and population structure, *u* is a vector of random genetic effects for each individual, **X** and **Z** are design matrices, and *e* is vector of residual effects. Variance matrix for *u* is 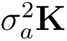 where 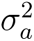 is the additive variance and **K** is the kinship matrix. Variance matrix for *e* is 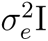, where 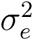 is the residual variance. For population structure correction, a principle component analysis was first done on the SNPs, and the components are used as fixed factors in the model. We chose the first four components for the analysis, as subsequent components explained very little of the variance (Figure S5). For testing association to TPC breadth, we used growth at either extreme temperature, 20 or 40 °C, and included the maximum growth rate at optimal temperature as an additional covariate to control for the variation in TPC elevation.

Testing for association single marker at a time has the disadvantage that nearby causal SNPs cannot be distinguished. It has been shown that testing for association iteratively always conditioning on the most significant marker improves statistical power (Segura *et al*., 2012). Therefore we used the BLINK method, which is a multilocus method that iteratively alternates between two different models as well as using LD information for the markers (Huang *et al*., 2018). Simulations have shown that the BLINK method has higher statistical power than single marker methods (Wang and Zhang, 2021; Huang *et al*., 2018) and it is computationally more efficient than the MLMM method of Segura *et al*. (2012) for large datasets (Wang and Zhang, 2021). Significance threshold for association was set to 0.01 after Bonferroni correction, which corresponds to a p-value of 6.78 *×* 10*^−^*^9^.

##### Allelic effects

For SNPs that were associated with growth at constant temperatures, we extracted their allelic effects from all temperatures, whether the associations were significant in those temperatures or not. For SNPs that were associated with TPC breadth, we examined their allelic effects in subsets of the nested association mapping population. We used a model that included the four principle components and maximum growth rate as covariates, effect of temperature, and interaction term with SNP and temperature. The model was:

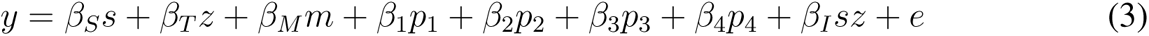

where *β_S_* is effect of the SNP, *β_T_*effect of temperature, *β_M_* effect of maximum growth rate, *β*_1_*…β*_4_ are effects of the population structure principle components, and *β_I_* the effect of interaction between SNP and temperature. We used predicted values to visualize the model results to account for the effects of population structure and the maximum growth rate.

## Results

### SNP density and linkage disequilibrium

To understand how the SNP density is related to patterns of LD, we evaluated the decay of linkage disequilibrium in our mapping population. First, we sampled pairs of SNPs randomly across all chromosomes to examine how LD decays over long distances across the chromosomes. We observed that on average LD does not persist over megabase scales (Figure S6).

Then we examined the decay of LD on a finer scale by sampling sets of consecutive SNPs evenly across the chromosomes. We observed that average LD decayed to very low levels after approximately 2 kb (Figure 1A). However, even if the average LD decayed fast, there was a large variation in LD, and some SNPs were in high LD over much longer distances (Figure 1A, S6). It is known that there is population structure between the populations from Louisiana and the Caribbean (Ellison *et al*., 2011), so high LD SNPs likely represent SNPs with allele frequency differences between the populations as well as recent mutations where recombination has not had enough time to break down LD.

**Figure 1:**
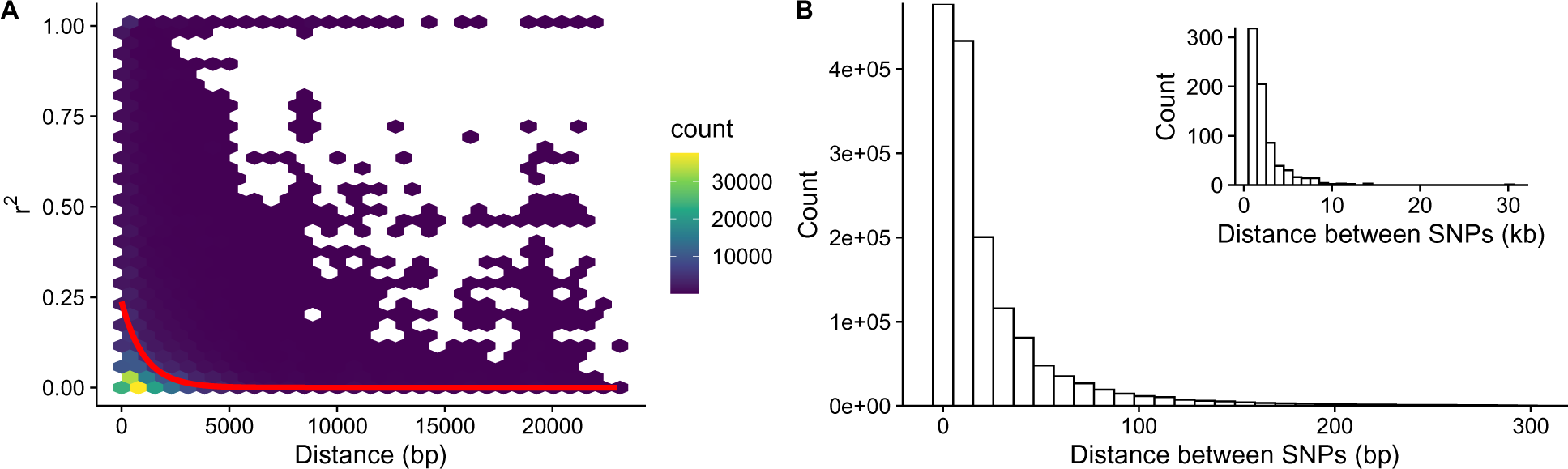
A) Linkage disequilibrium between 346 500 pairwise SNP combinations sampled evenly across the seven chromosomes. Only strains belonging to the set of natural populations are included in the analysis. In the plot data has been binned into hexes. Red line shows exponential decay fit. B) Distribution of physical distance between consecutive SNPs in the natural populations. Inset shows SNPs that are further than 1 kb apart, note the different scales of the two histograms.

When we compared the distribution of physical distances between consecutive SNPs we observed that 99% of SNPs are within 215 bp or less of the next SNP (Figure 1B). Only few SNPs had distances longer than 1 kb to the next SNP (Figure 1B), and these were mostly SNPs near the centromeric regions. Since LD decays over longer distances than the SNP density in our mapping population, we should have a good chance to detect associations, even if our re-sequencing has for some reason missed a causal SNP. Yet, our results suggest that association studies in *N. crassa* need a very dense genotyping to have a chance to detect all causal variants, and long distance haplotype tagging cannot be relied on.

### Genome-wide association

When testing for associations at each temperature, we observed 12 associations that were significant after Bonferroni correction at the 0.01 level across all temperatures (Figure 2). We observed both unique associations in a given temperature and associations that were shared across multiple temperatures (Figure 2, Table 2).

**Figure 2:**
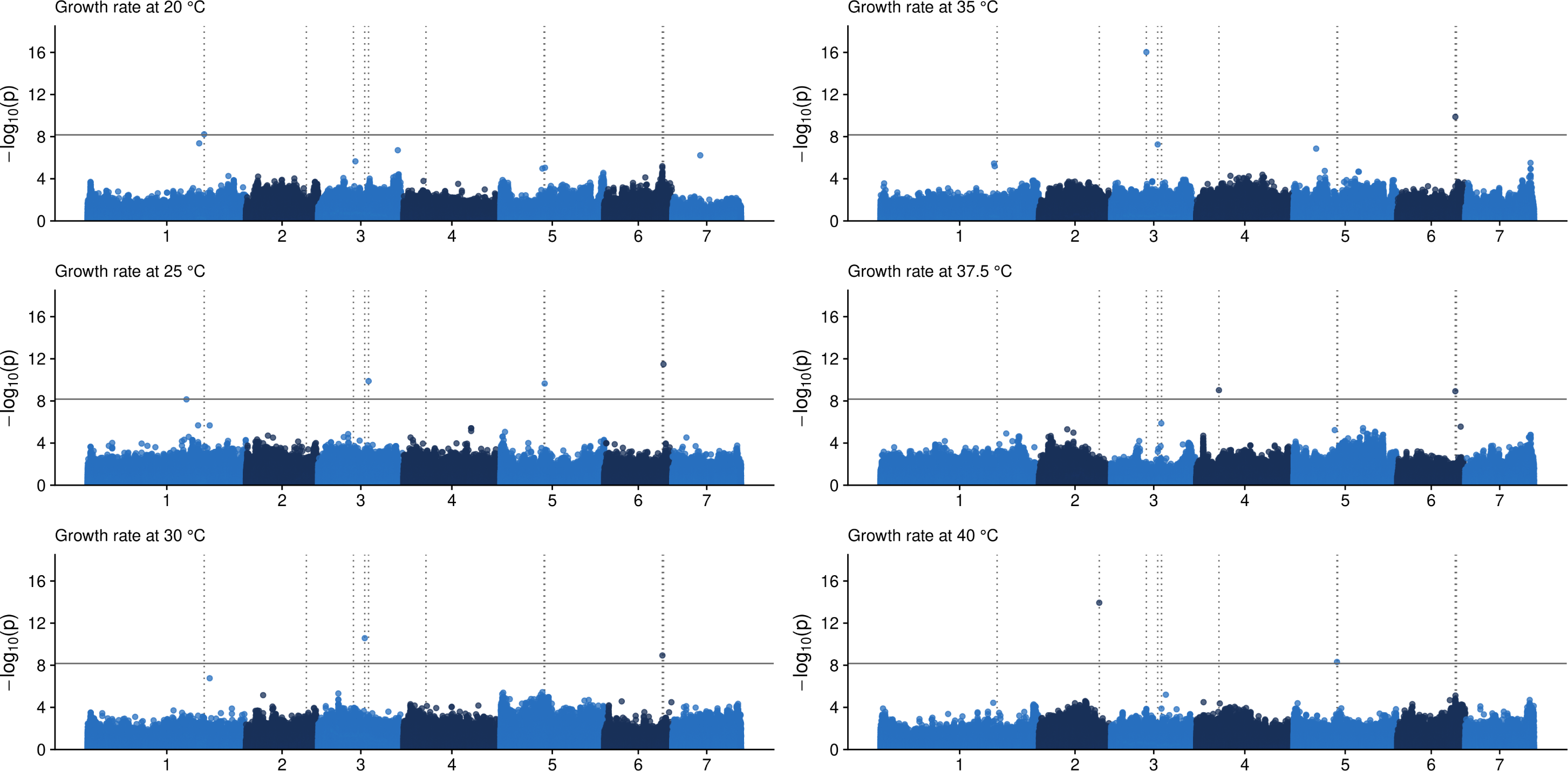
Manhattan plots for growth rate at different temperatures. Solid horizontal line denotes p-value threshold of 0.01 after Bonferroni correction. Dotted vertical lines highlight positions of SNPs that were significant in at least one of the temperatures. Note that because we used a multi marker method to test for associations, we do not expect to see a "skyscraper" of significant SNPs around a significant SNP in the Manhattan plot.

**Table 2:** SNPs that were associated with growth. Temp. = Temperature, Chr. = Chromosome. For intergenic SNPs the nearest gene is reported, and distance to the nearest gene is given in the variant column.

| Temp. (°C) | Chr. | Position (bp) | p-value | Gene | Annotation | Variant |
| --- | --- | --- | --- | --- | --- | --- |
| 20 | 1 | 7 214 188 | $6.00 \times 10^{-9}$ | NCU00775 | isocitrate dehydrogenase subunit 1, *tca-6* | intergenic (downstream 156 bp) |
| 25 | 3 | 3 119 653 | $1.31 \times 10^{-10}$ | NCU00236 | flavoprotein oxygenase, *fpo-1* | synonymous |
| 25 | 5 | 2 747 660 | $2.22 \times 10^{-10}$ | NCU01043 | hypothetical protein | 3'UTR |
| 25 | 6 | 3 669 489 | $3.27 \times 10^{-12}$ | NCU17138 | haloacid dehalogenase like protein | missense (S/P) |
| 30 | 3 | 2 881 603 | $2.69 \times 10^{-11}$ | NCU00170 | DNAJ domain-containing protein | intergenic (upstream 512 bp) |
| 30 | 6 | 3 600 716 | $1.18 \times 10^{-9}$ | NCU05137 | non-anchored cell wall protein 1, *ncw-1* | synonymous |
| 35 | 3 | 2 179 382 | $9.39 \times 10^{-17}$ | NCU06278 | elongation factor 2, variant 2, *rbg-37* | missense (G/S) |
| 35 | 6 | 3 600 716 | $1.32 \times 10^{-10}$ | NCU05137 | non-anchored cell wall protein 1, *ncw-1* | synonymous |
| 37.5 | 4 | 1 399 273 | $9.52 \times 10^{-10}$ | NCU07904 | short chain dehydrogenase/reductase | intronic, splice polypyrimidine tract |
| 37.5 | 6 | 3 599 026 | $1.20 \times 10^{-9}$ | NCU05138 | glycosyltransferase family 25 protein, *gt25-1* | missense (E/Q) |
| 40 | 2 | 3 743 231 | $1.17 \times 10^{-14}$ | NCU08705 | NACHT domain-containing protein | intergenic (upstream 1182 bp) |
| 40 | 5 | 2 708 042 | $4.85 \times 10^{-9}$ | NCU01024 | MICOS complex subunit MIC12 | missense (L/I) |

There were six non-coding variants among the significant associations, these included two synonymous variants, one 3’UTR, three intergenic variants that were close to genes, and one intronic variant (Table 2). If these non-coding variants are true positives, then they are presumably regulatory variants.

Four of the associations were missense variants, these included a missense variant in the gene *NCU17138*, encoding haloacid dehalogenase (HAD) like protein (Table 2), which was associated with growth rate at 25 °C. A missense variant in the gene *rbg-37*, encoding elongation factor 2 variant 2, that was associated in 35 °C. There was also a missense variant in the gene *gt25-1*, encoding glycosyltransferase family 25 protein, that was associated with growth at 37.5 °C, and which was within 1690 bp of the shared association between 30 °C and 35 °C (Table 2). This raises the possibility that there is one true association within this region. Furthermore, a missense variant in the gene *NCU01024*, encoding MICOS complex subunit MIC12, was associated at 40 °C.

To evaluate whether the associations we observed were true positives, we examined the segregation of associated variants among the strains sampled from natural populations and the different families. In general, we observed that whenever a significant SNP segregated in multiple families its effects were congruent across different families and the natural populations (Figure S7, S8, S9, S10, S11, and S12). In some cases, the directions of the allelic effects on growth rate were the same but the effects were of different magnitude (e.g. Figure S7), one explanation for this is epistasis, so that the magnitude of allelic effects varies in different families because of the genetic background in which they occur. While segregation in different parts of the nested association mapping population does not constitute a validation of the SNPs in an independent set of data, it does seem unlikely that a SNP which shows an effect in the same direction in three different families and in the natural populations, such as the SNP in chromosome 5: 2 708 042 (Figure S12), would be a false positive. We further tested if we could detect which genes influence thermal performance curve shape, by performing GWAS for optimum temperature and TPC breadth, where variation in elevation has been taken into account. We observed one synonymous association for TPC optimum temperature in the gene *stk-32* encoding protein kinase domain-containing protein ppk32 (Table 3). We found seven associations for growth at 20 °C conditioned on maximum growth rate. Three of these SNPs were synonymous variants, first in the gene *NCU02025* encoding Nup53p nucleoporin-like protein, second in the gene *ptk-2* encoding a non-specific serine/threonine protein kinase, and third in the gene *rbg-11* encoding periodic tryptophan protein 2 homolog. At 40 °C conditioned on maximum growth rate we found three associations that were intergenic polymorphisms. None of the associations were shared among the three traits (Figure 3). There was no overlap between significant associations for growth rate at different temperatures and the TPC shape traits (Table 2 and 3). All of the SNPs associated with TPC shape traits were potential regulatory variants and no missense variants were associated with TPC shape (Table 3). Some of them, such as SNP at 1: 6 331 236 and SNP at 3: 3 616 832 that were associated with growth at 40 °C conditioned for maximum growth rate, were rather far away from the nearest genes (Table 3).

**Figure 3:**
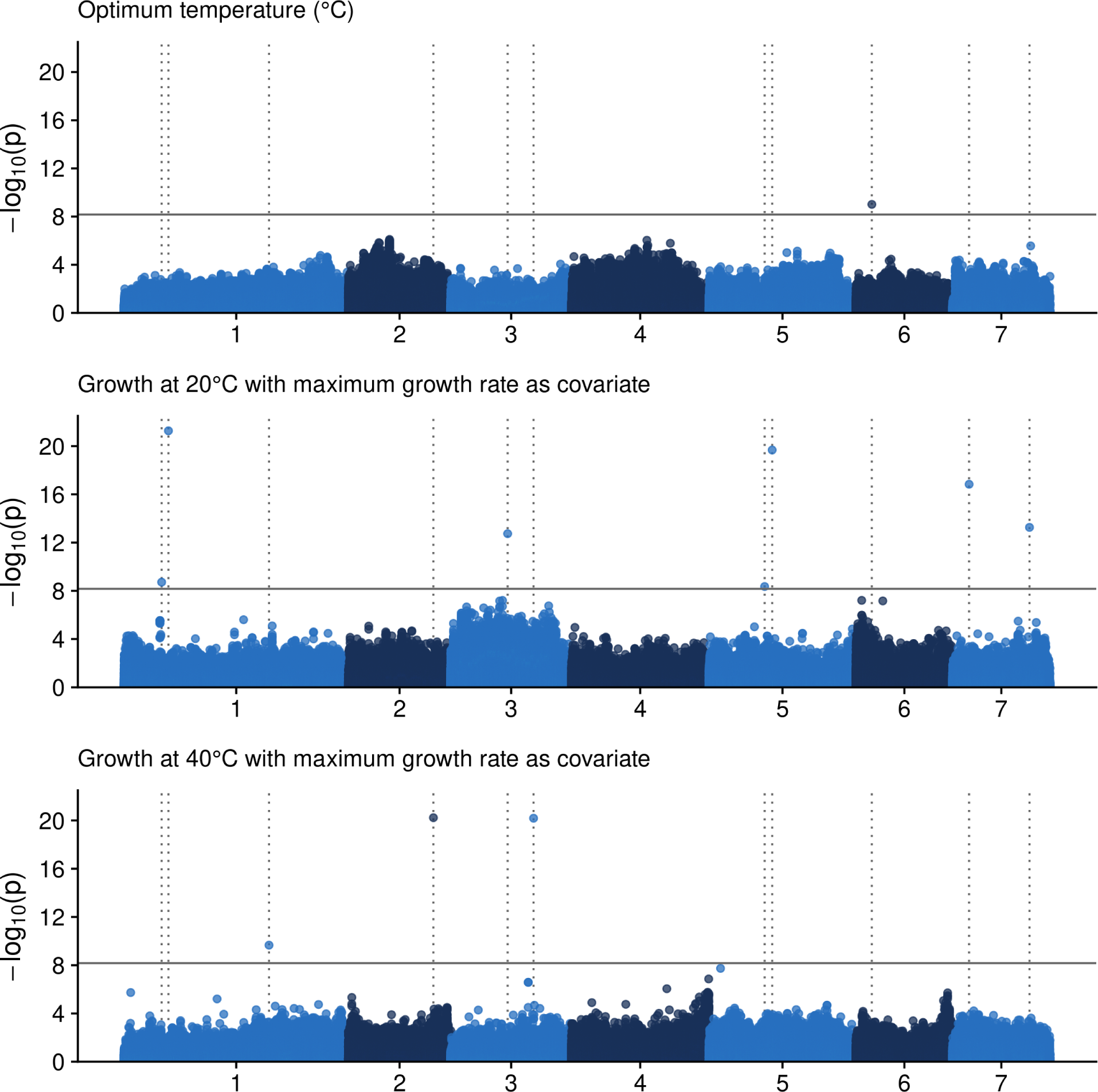
Manhattan plots for thermal performance curve traits. Solid horizontal line denotes p-value threshold of 0.01 after Bonferroni correction. Dotted vertical lines highlight positions of SNPs that were significant for at least one trait.

**Table 3:**
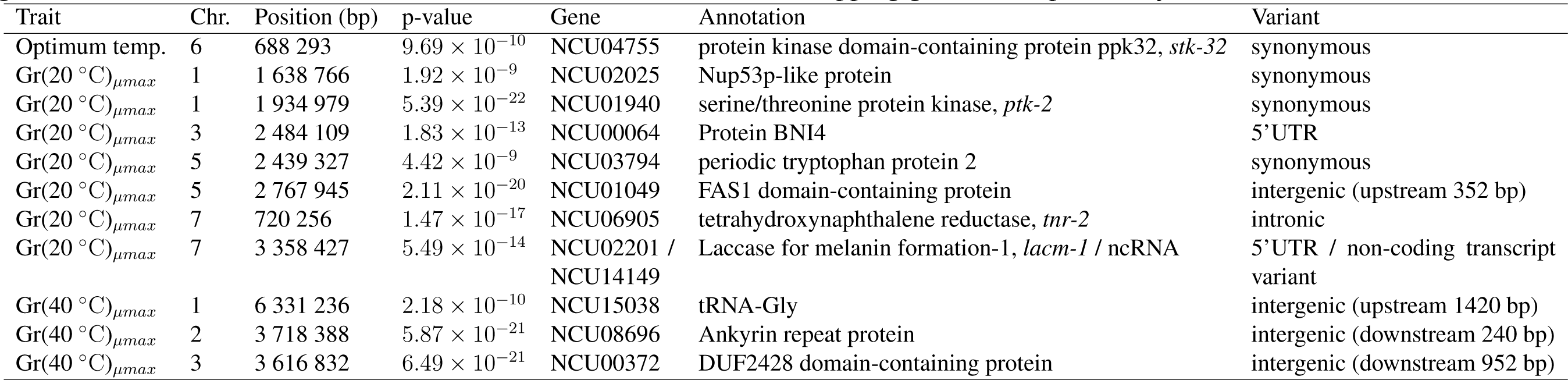
SNPs that were associated with TPC shape. Chr. = Chromosome. For intergenic SNPs the nearest gene is reported, and distance to the nearest gene is given in the variant column. For the trait column, Gr(X)*_µmax_* are growth rates at temperature X where maximum growth rate was used as a covariate in the model. Annotations for overlapping genes are separated by "/".

| Trait | Chr. | Position (bp) | p-value | Gene | Annotation | Variant |
| --- | --- | --- | --- | --- | --- | --- |
| Optimum temp. | 6 | 688 293 | $9.69 \times 10^{-10}$ | NCU04755 | protein kinase domain-containing protein ppk32, *stk-32* | synonymous |
| $Gr(20^\circ C)_{\mu max}$ | 1 | 1 638 766 | $1.92 \times 10^{-9}$ | NCU02025 | Nup53p-like protein | synonymous |
| $Gr(20^\circ C)_{\mu max}$ | 1 | 1 934 979 | $5.39 \times 10^{-22}$ | NCU01940 | serine/threonine protein kinase, *ptk-2* | synonymous |
| $Gr(20^\circ C)_{\mu max}$ | 3 | 2 484 109 | $1.83 \times 10^{-13}$ | NCU00064 | Protein BNI4 | 5'UTR |
| $Gr(20^\circ C)_{\mu max}$ | 5 | 2 439 327 | $4.42 \times 10^{-9}$ | NCU03794 | periodic tryptophan protein 2 | synonymous |
| $Gr(20^\circ C)_{\mu max}$ | 5 | 2 767 945 | $2.11 \times 10^{-20}$ | NCU01049 | FAS1 domain-containing protein | intergenic (upstream 352 bp) |
| $Gr(20^\circ C)_{\mu max}$ | 7 | 720 256 | $1.47 \times 10^{-17}$ | NCU06905 | tetrahydroxynaphthalene reductase, *tnr-2* | intronic |
| $Gr(20^\circ C)_{\mu max}$ | 7 | 3 358 427 | $5.49 \times 10^{-14}$ | NCU02201 /<br>NCU14149 | Laccase for melanin formation-1, *lacm-1* / ncRNA | 5'UTR / non-coding transcript<br>variant |
| $Gr(40^\circ C)_{\mu max}$ | 1 | 6 331 236 | $2.18 \times 10^{-10}$ | NCU15038 | tRNA-Gly | intergenic (upstream 1420 bp) |
| $Gr(40^\circ C)_{\mu max}$ | 2 | 3 718 388 | $5.87 \times 10^{-21}$ | NCU08696 | Ankyrin repeat protein | intergenic (downstream 240 bp) |
| $Gr(40^\circ C)_{\mu max}$ | 3 | 3 616 832 | $6.49 \times 10^{-21}$ | NCU00372 | DUF2428 domain-containing protein | intergenic (downstream 952 bp) |

### Allelic effects

When we plotted the allelic effects of all SNPs that had a significant association in one of the temperatures across all temperatures, we observed that in general minor alleles decreased growth (Figure 4). Some SNPs had an effect in only one temperature, such as the SNP in chromosome 2: 3 743 231 for which the minor allele decreased growth rate at 40 °C but had no effect in other temperatures (Figure 4). Similarly for the SNP in chromosome 5: 2 708 042 for which minor allele decreased growth rate only at 40 °C. Some SNPs had a tendency to have largest effects at either low or high temperatures, such as the SNP in chromosome 1: 7 214 188 and the SNP in chromosome 6: 3 599 026 (Figure 4). One SNP at chromosome 3: 2 881 603 had the largest effects at the optimal temperature of 35 °C and its effect decreased when moving away from the optimum temperature, which suggest the possibility of thermodynamic scaling (Figure 4).

**Figure 4:**
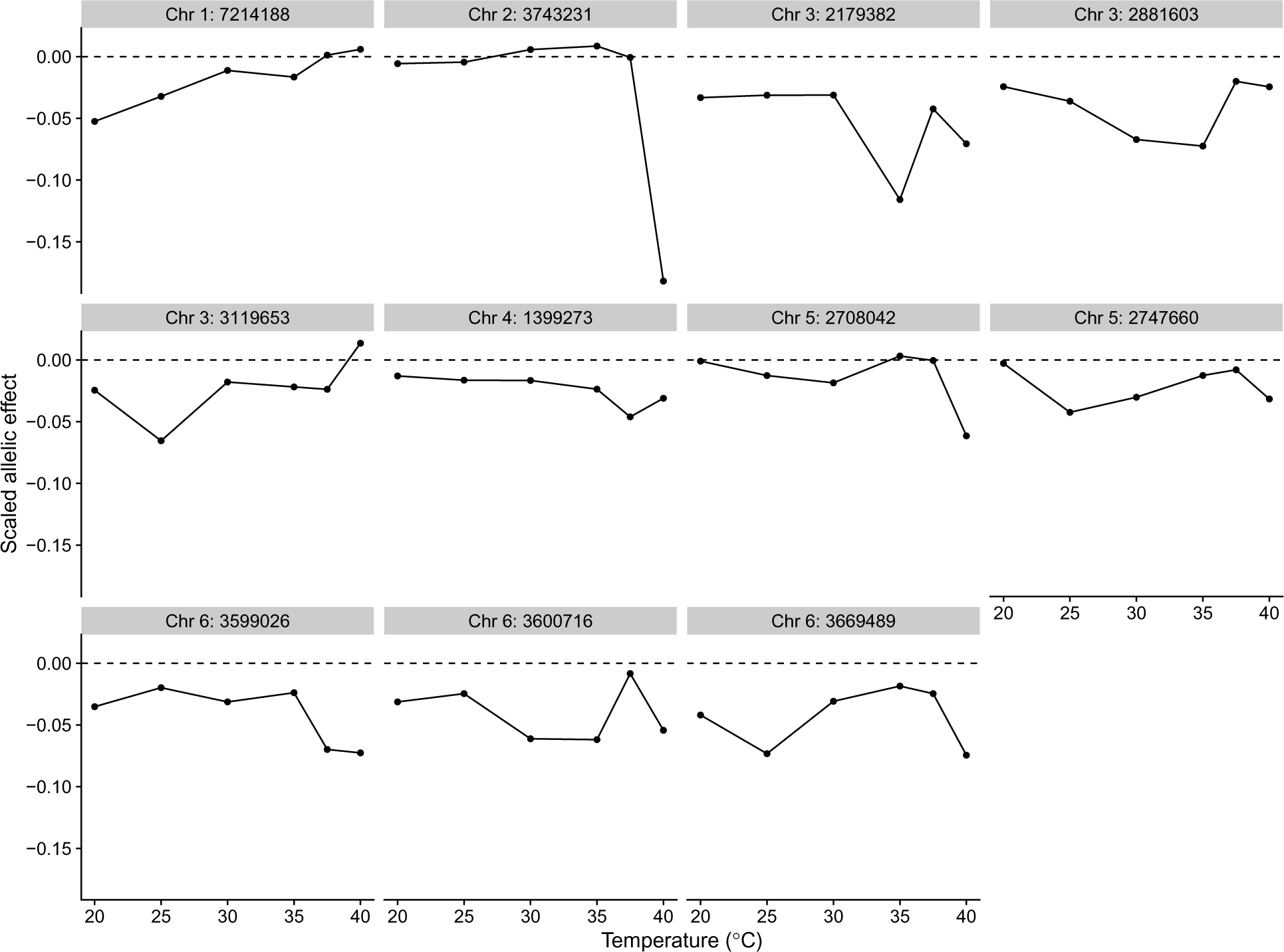
Scaled allelic effects of SNPs that were associated with growth at any temperature over all of the measured temperatures. Allelic effects are reported relative to the major allele, that is, if the minor allele decreases growth rate, the allelic effect has a negative sign.

Then we examined the relationship between the size of the allelic effects and minor allele frequencies in the natural populations. We observed a negative relationship between the absolute scaled allelic effect and the minor allele frequency in the natural populations (Figure 5). Since we observed that for all significant associations the minor allele decreased growth, and that allelic effects get smaller as allele frequency increases in the natural populations, these associated alleles are likely to be mildly deleterious alleles segregating at low frequencies in the natural populations.

**Figure 5:**
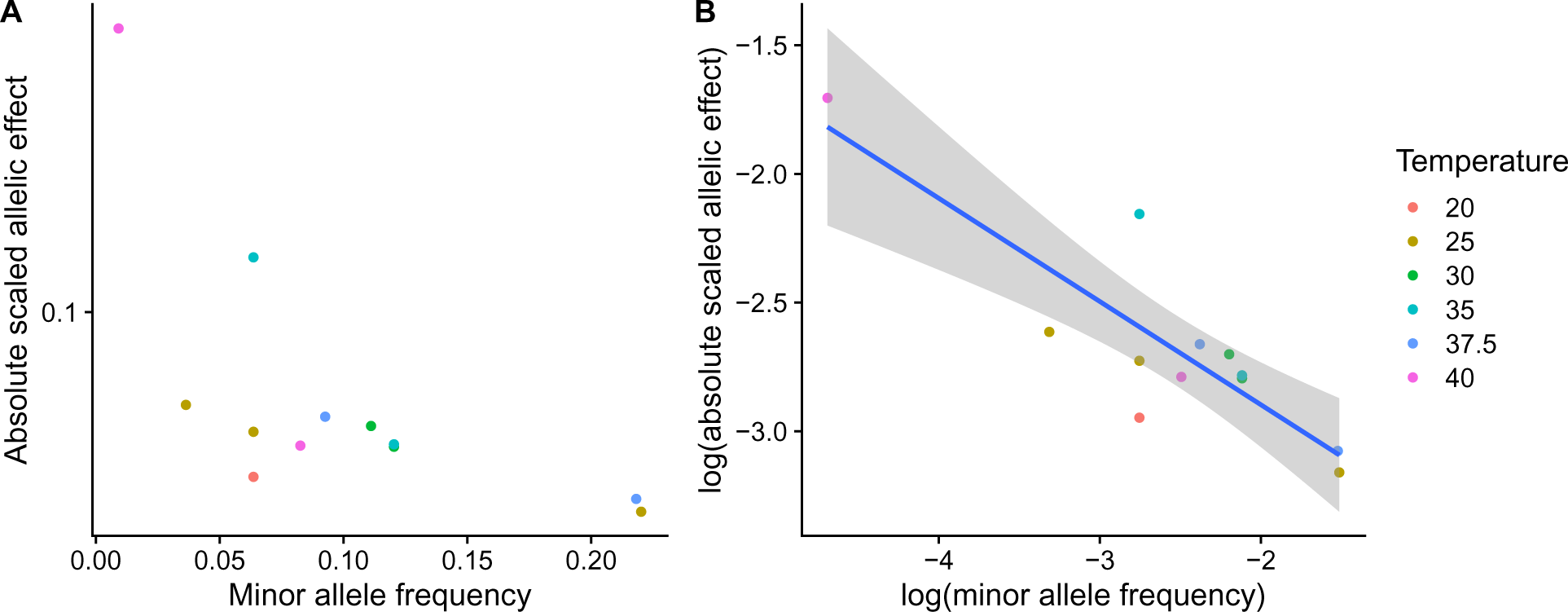
A) Absolute scaled allelic effects of associated SNPs plotted against minor allele frequency in natural populations. Minor allele frequencies are frequencies among the strains from the natural populations, not in the whole nested association mapping population. B) The same data in a log-log plot, allelic effects get smaller as minor allele frequency increases.

Then we examined the allelic effects of SNPs that were associated with growth at either 20 or 40 °C when maximum growth rate was used as a covariate. Since the effect of variation in elevation was removed from this data, we were interested if there were any trade-offs between the temperatures. That is, did the allelic effect switch its sign at the other temperature, regardless of genome-wide significance. Out of the ten SNPs that were significantly associated with TPC breadth in either direction, for five SNPs we observed some indications of trade-offs (Figure 6).

**Figure 6:**
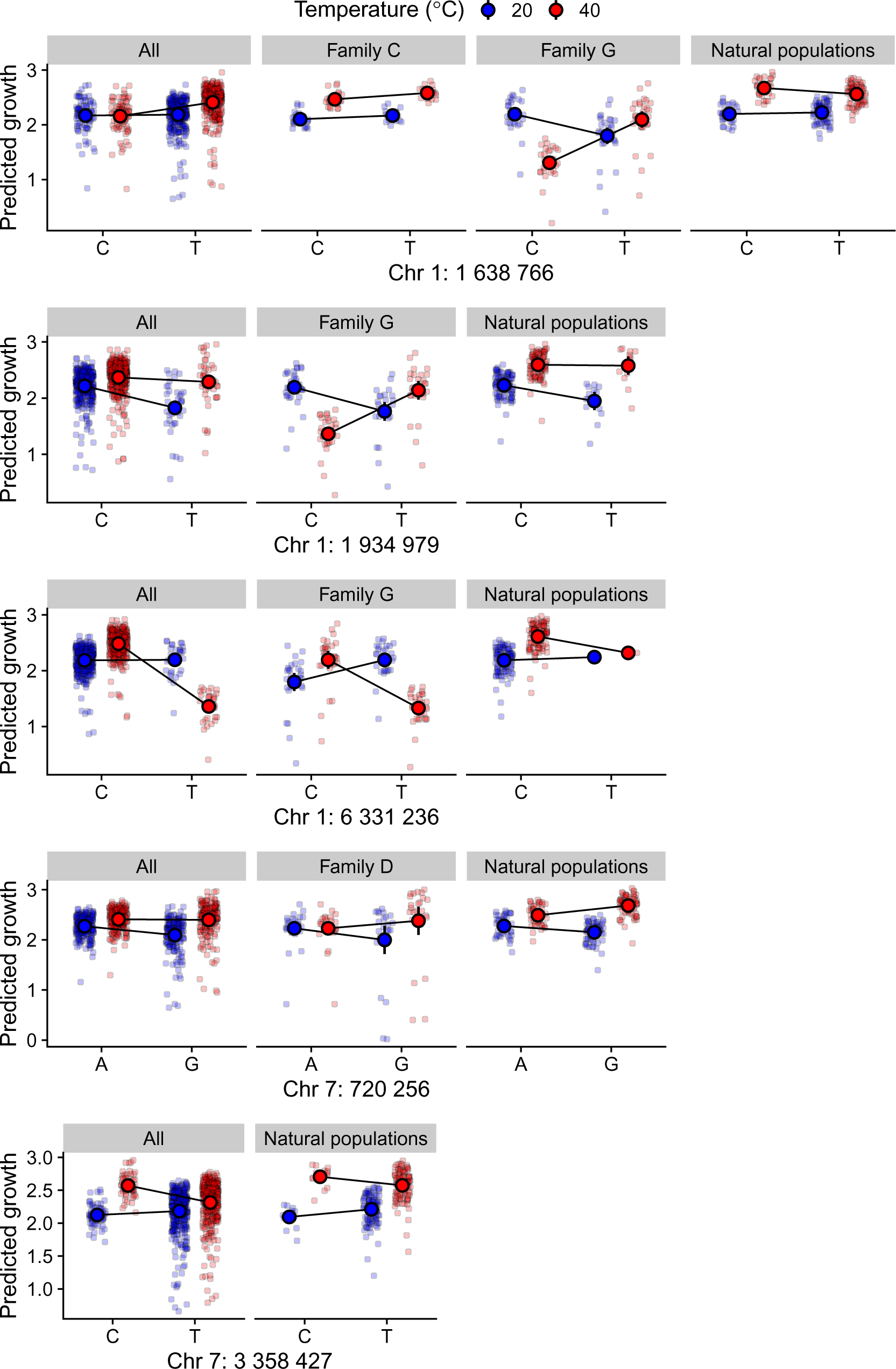
Allelic effects of SNPs associated with TPC breadth, which show potential trade-offs between 20 and 40 °C. Y-axis shows model predictions.

Three SNPs in chromosome 1 displayed a potential trade-off. There was a trade-off for the SNP in chromosome 1 at position 1 638 766 among the family G strains. Effect of the SNP at 20 °C was *−*0.39, *p* = 0.01, and the effect at 40 °C was 0.79, and the interaction term had a *p* = 1.68 *×* 10*^−^*^11^. However, no trade-off was observed among strains belonging to family C or among the strains from natural populations, where this SNP also segragated. There was also a nearby SNP in chromosome 1 at position 1 934 979 where a trade-off was also observed among strains of family G, the effect of the SNP at 20 °C was *−*0.43, *p* = 0.01, and the effect of the SNP at 40 °C was 0.77, and *p* = 3.20 *×* 10*^−^*^12^ for the interaction term. But this trade-off was not observed among the natural strains, where the SNP had an effect only at 20 °C (Figure 6). These two SNPs are close together physically, and within family G, these effects are likely similar due to linkage. So we cannot be certain which of these SNPs is causally behind the trade-off, or whether the observed trade-off in family G is the joint effect of multiple linked loci. We also observed that for a SNP in chromosome 1 at position 6 331 236 estimates of allelic effects had opposite signs for family G, but the effect of SNP was not significant at 20 °C, *p* = 0.33 even if the interaction term was significant, *p* = 8.59 *×* 10*^−^*^15^. We could not assess this potential trade-off in other parts of the nested association mapping population, since this SNP is carried only by strain 851, which is one of the parents of family G. For these three SNPs the trade-off is present potentially only in family G, and we could not confirm it in any other part of the mapping population. Therefore, we view the evidence for trade-offs as weak for these SNPs.

Another two SNPs in chromosome 7 also showed a potential trade-off. For the SNP in chromosome 7 at position 720 256 among family D had opposite signs for allelic effects: *−*0.23 at 20 °C, *p* = 0.11 and 0.15 at 40 °C, *p* = 0.0003 for the interaction term. Since the main effect of the SNP was not significant this remains only suggestive. However, for the natural strains the estimate of the SNP effect at 20 °C was *−*0.13, *p* = 0.02 and effect at 40 °C was 0.19, and *p* = 0.0001 for the interaction term. So, a significant trade-off was present among the natural strains for this SNP, and the effects go in the same direction in family D. The second SNP in chromosome 7 at position 3 358 427 had a significant trade-off among the natural strains. The effect of the SNP at 20 °C was 0.12, *p* = 0.003 and the effect at 40 °C was *−*0.14, *p* = 0.03 for the interaction term. Observing a significant trade-off for the natural populations seems stronger evidence than just for a single family.

The other SNPs did not have significant trade-offs (Figure S13). Either the allelic effects on growth had the same direction at both temperatures, or the effect at the other temperature was weak, or there was an effect only at the other temperature.

## Discussion

While association mapping has been previously used in *N. crassa* to discover genes affecting quantitative traits (Palma-Guerrero *et al*., 2013), our results demonstrate that our nested association mapping population can be successfully used to map loci underlying quantitative traits. As *N. crassa* strains can be stored in the freezer, it is possible to use this population again to investigate other traits. In the future, the statistical power could be improved by making more crosses using more parents, so that singleton SNPs among the natural strains will be segregating in multiple families. This would also improve the estimation of allelic effects if the alleles occur in multiple backgrounds.

Our results allow some insight into the identity of the loci that influence growth at different temperatures and the functions of those loci. Many of the genes associated with growth at different temperatures seem to be genes important for growth, rather than temperature response specifically.

For example, the gene *tca-6* associated at 20 °C functions in the tricarboxylic acid cycle to release energy from stored nutrients. The gene *fpo-1* associated at 25 °C encodes for flavoprotein oxygenase, which belongs to a class of enzymes that function, e.g., in energy production, degradation of organic material, biosynthesis of primary metabolites like pigments (Kerschbaumer *et al*., 2022), and amino acid metabolism (Huberman *et al*., 2021). Deletion mutants of *fpo-1* gene have been shown to suffer from reduced reproductive ability (Lan *et al*., 2021). Moreover, the gene *ncw-1*, which was associated in both 30 and 35 °C, is known to function in carbohydrate metabolic process, as its deletion promotes cellulase production by increasing cellobiose uptake in *N. crassa* through transporter expression (Lin *et al*., 2017).

We did observe one locus that had a clear link to temperature response. The gene *NCU00170* associated at 30 °C encodes DnaJ domain-containing protein. DnaJ is an important molecular chaperone and a member of the Hsp40 family, which regulates the activity of Hsp70s and fungal ATPase machinery (Roy and Tamuli, 2022). Heat shock proteins (Hsps) are involved in a wide array of biological processes in fungi, including growth, development, and general stress resistance (Abu Bakar *et al*., 2020). In *Neurospora*, DnaJs are the largest class of Hsps, proteins which are produced to prevent protein unfolding and denaturation under sub-optimal or stressful conditions (Borkovich *et al*., 2004; Kapoor and Roy, 2014).

Further genes were also related to growth. At 35 °C we found a missense SNP in the gene *rbg-37* encoding a variant elongation factor 2. This protein binds the ribosomal subunits together into a ribosome and functions in the chain elongation during protein synthesis. Previously the gene *cot-3* encoding translation elongation factor 2 has been found to be involved in hyphal growth at elevated temperatures in *N. crassa* (Propheta *et al*., 2001). We also found a missense SNP associated with growth at 37.5 °C in the gene *gt25-1* encoding glycosyltransferase family 25 protein. Glycosyltransferases have been linked to fungal hyphal growth on solid matrices (King *et al*., 2017). Finally, a missense SNP in the gene *NCU01024* was associated at 40 °C, this gene encodes the MICOS complex subunit MIC12, which is part of the mitochondrial inner membrane.

For the TPC shape traits, we observed one association with the TPC optimum temperature itself, for the gene *stk-32*, which encodes a protein kinase. In filamentous fungi this gene has been linked to the regulatory mechanism of mycelial growth under heat stress (Yang *et al*., 2022), and the deletion mutants of the *stk-32* gene in *N. crassa* have reduced hyphal growth (Park *et al*., 2011). When we tested for associations with TPC breadth, and used the maximal growth rate as a covariate to remove the effect of elevation variation, we observed that many of the associated genes had growth and cell wall related functions. At 20 °C there was as association for the gene *ptk-2* encoding a protein kinase that in *N. crassa* is involved in the hyphal tip-growth (Lew and Kapishon, 2009). Another association at 20 °C was in the gene *NCU00064* encoding the BNI4 protein which is known to affect the cell wall building and hyphal growth in *N. crassa* (Verdín *et al*., 2019). We observed also an association in the gene *rbg-11*, which encodes periodic tryptophan protein 2 homolog that affects cell polarity and growth. In yeast, this protein is involved in ribosome biogenesis and regulation of the cell cycle (Neer *et al*., 1994).

In addition to these genes involved in growth processes, we found two genes involved in melanin biosynthesis to be associated with TPC breadth. The gene *tnr-2* encodes a tetrahydroxynaphthalene reductase, which is an enzyme that functions in the fungal biosynthesis of melanin. Melanin strengthens the cell wall and is important for adaptation to stressful conditions (Suthar *et al*., 2023). The other gene involved in melanin production was *lacm-1*. This gene encodes an enzyme that catalyzes melanin formation for making strong cell wall structure, also in perithecium and ascospores (Ao *et al*., 2019). Furthermore, this gene overlaps with a non-coding RNA gene *NCU14149*, but no obvious function can be assigned to this ncRNA.

For genes associated with TPC breadth at 40 °C, the gene *NCU15038* encodes a glycine transfer RNA. Glycine transfer RNA can modify fungal cell walls and membrane constituents by adding amino acids to them (Grob *et al*., 2022). Another associated gene was *NCU08696*, which encodes an ankyrin repeat protein that functions in ion transmembrane transporter activity and protein–protein interactions involved in the formation of transcription complexes. Mutations in genes that encode ankyrin-like proteins can cause defects in gene expression, and it has been linked to mycelial growth in fungi (Situ *et al*., 2023). The final associated gene *NCU00372*, was a DUF2428 domain-containing protein, this protein is highly conserved and known to be required for tRNA methylation in yeasts and interestingly, is in general involved in energy metabolism and growth response during cold conditions (Dong *et al*., 2019).

We know that for *N. crassa* the majority of genetic variation for growth at different temperatures is in TPC elevation (Moghadam *et al*., 2020). When we tested for associations at specific temperatures, we observed both unique associations, and associations that were shared across multiple temperatures. Growth in nearby temperatures has a positive and high genetic correlation (Moghadam *et al*., 2020), so shared associations are expected. It is perhaps surprising that so few shared associations were detected. Since we can only detect associations with high and moderate effects, it may be that alleles that affect growth at multiple temperatures have smaller effects in general and we miss many of the shared associations. Moreover, there was no strong evidence of thermodynamic scaling for allelic efffects, as we observed only one locus for which allelic effects were compatible with thermodynamic scaling. While there is a correlation between optimum temperature and maximum growth rate at the phenotypic level (Moghadam *et al*., 2020), it seems that allelic effects do not often scale this way. Rather allelic effects tend to be limited for certain temperature ranges.

For growth at specific temperatures we observed that effects of the minor allele had a negative correlation with minor allele frequency in the natural populations. This suggests that alleles with large effects on growth are mostly deleterious, and are being kept at low frequencies by negative selection (Barton and Keightley, 2002; Keightley and Lynch, 2003).

Allelic variation with antagonistic pleiotropy can lead to divergent adaptation when certain alleles act to increase fitness in one environment and conversely decrease in others, which is a pre-requisite for trade-offs in TPCs (Angilletta *et al*., 2003; Yamahira *et al*., 2007; Berger *et al*., 2013; Murren *et al*., 2015). In a study of hot and cold adapted experimental lines of dung fly (*Sepsis punctum*), temperature-specific allelic effects on growth rate were found after 20 generations, but no trade-offs in TPC shape were observed (Berger *et al*., 2014). Another study with European grayling (*Thymallus thymallus*) adapted to differing thermal environments demonstrated temperature-driven gene expression changes constrained by the level of gene pleiotropy (Papakostas *et al*., 2014). Interestingly, they found that genes with low pleiotropy levels were the main drivers of the changes in gene expression, whereas highly pleiotropic genes had limited expression response to temperature treatment.

Antagonistic pleiotropy can be produced for example by alleles that encode different enzyme variants acting differently across temperatures (Huey and Kingsolver, 1989). However, typically the genetic variance for temperature specificity of enzymatic reactions is low (Hochachka and Somero, 2002; Latimer *et al*., 2014). Other possible mechanisms are the frequency change at loci affecting overall performance but with differing strength between temperatures, and the difference in the magnitude of the allelic effects, for example, between cold and hot temperatures (Latimer *et al*., 2015).

We only detected weak evidence for trade-offs out of the ten SNPs associated with TPC breadth at either hot or cold temperatures. Potential trade-offs were detected for five SNPs, and for three of those the evidence of trade-off came from family G, and the trade-off was not supported by the strains from natural populations. It may be that there is a modifier locus present in family G, that is an epistatic effect, or that effects of multiple loci that affect growth only at hot or cold temperature are joined together due to linkage and appear as a trade-off. Either way, evidence of trade-off seems relatively weak for these three SNPs. For the two remaining SNPs evidence of trade-off was stronger, since the trade-off was observed also among the natural populations.

Since we know that at the phenotypic level this hot-cold trade-off represents a minor aspect of the genetic variation (Moghadam *et al*., 2020), it is not that surprising that only few loci display evidence of trade-offs. Our results highlight the need to reconsider some aspects of theories related to TPC evolution, where antagonistic pleiotropy is predicted to have an important role in maintaining variation in TPCs (Lynch and Gabriel, 1987; Gilchrist, 1995). Our results suggest that some trade-offs at the allelic level exist, but it seems that in majority of cases alleles have temperature specific effects. Our results are in line with a genome-wide study of thermal tolerance in *Drosophila melanogaster*, while some genes showed pleiotropic effects shaping tolerance at upper and lower thermal limits (Lecheta *et al*., 2020), the heat and cold tolerance had no overlapping SNPs, so distinct molecular processes evolving indepently in response to changing environmental conditions are more likely.

## Conclusion

In summary, we identified that many loci contributing to TPC variation are loci that may affect growth and metabolism in general, as well as the fungal cell wall. Finding genes related to growth is understandable, since most of this variation is related to TPC elevation, but the associated loci have temperature specific alleles. We did detect some loci that had been previously linked with temperature stress or stress response in general. We also found that trade-offs between hot and cold temperatures do exist at the allelic level, but they are a minor part of the overall variation. Therefore, variation in TPCs is unlikely to be driven solely by trade-offs.

### Data access

The genome sequencing data generated in this project has been deposited to Sequence Read Archive under accession number XXXXX-XXXX, and RAD-sequencing data under accession number PRJNA1109407. Genotypes are available on Zenodo: https://doi.org/10.5281/zenodo.11120317, and related scripts for the nested association mapping population can be accessed at: https://github.com/ikron/Neurospora_NAM_population. Phenotypes and analysis scripts can be accessed at: https://github.com/ikron/TPC_GWAS.

## Acknowledgements

The authors wish to acknowledge CSC – IT Center for Science, Finland, for computational resources, and the Fungal Genetics Stock Center for the strains used in this study. This study was funded by grants from Emil Aaltonen foundation to IK and ER, University of Jyväskylä Doctoral Programme in Biological and Environmental Science to ER, Ella & Georg Ehrnrooth foundation to IK, and Academy of Finland Research Fellowships to IK (no. 321584) and TK (no. 278751).

## Author contributions

The study was conceived by I.K., T.K, and E.R. Experiments were performed by N.N.M., H.-R.L., K.S., P.A.M.S., E.R. and I.K. Data was analyzed by I.K. and I.K. wrote the manuscript with support from E.R.. All authors edited the final manuscript.

## Supplementary Information

### Supplementary Figures

**Figure S1:**
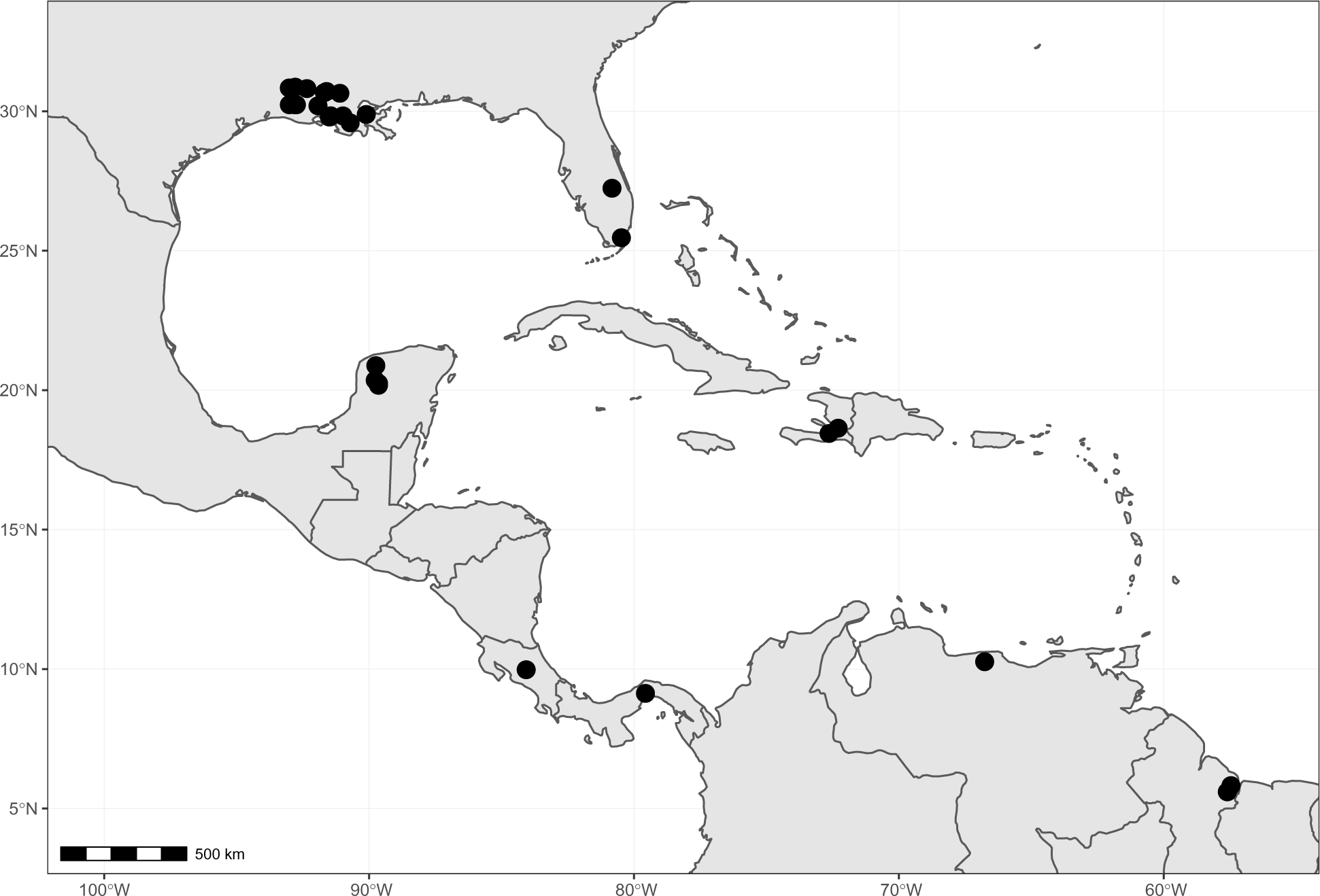
Sampling locations around Southeastern United States and the Caribbean, where the strains originating from nature have been collected.

**Figure S2:**
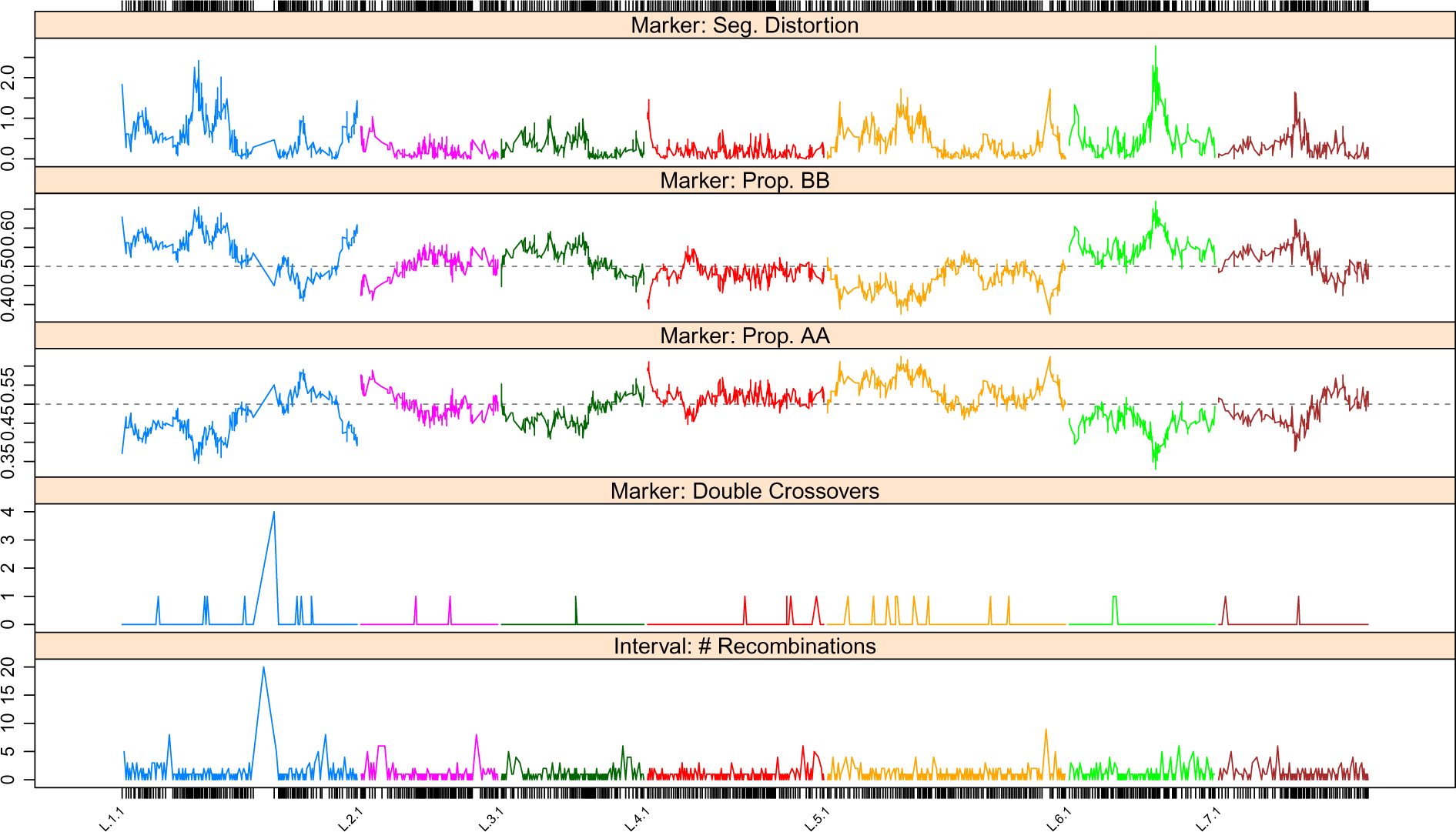
An example of segregation distortion, marker frequencies, doubles crossovers and recombination events in family A offspring. Some double crossover events are observed.

**Figure S3:**
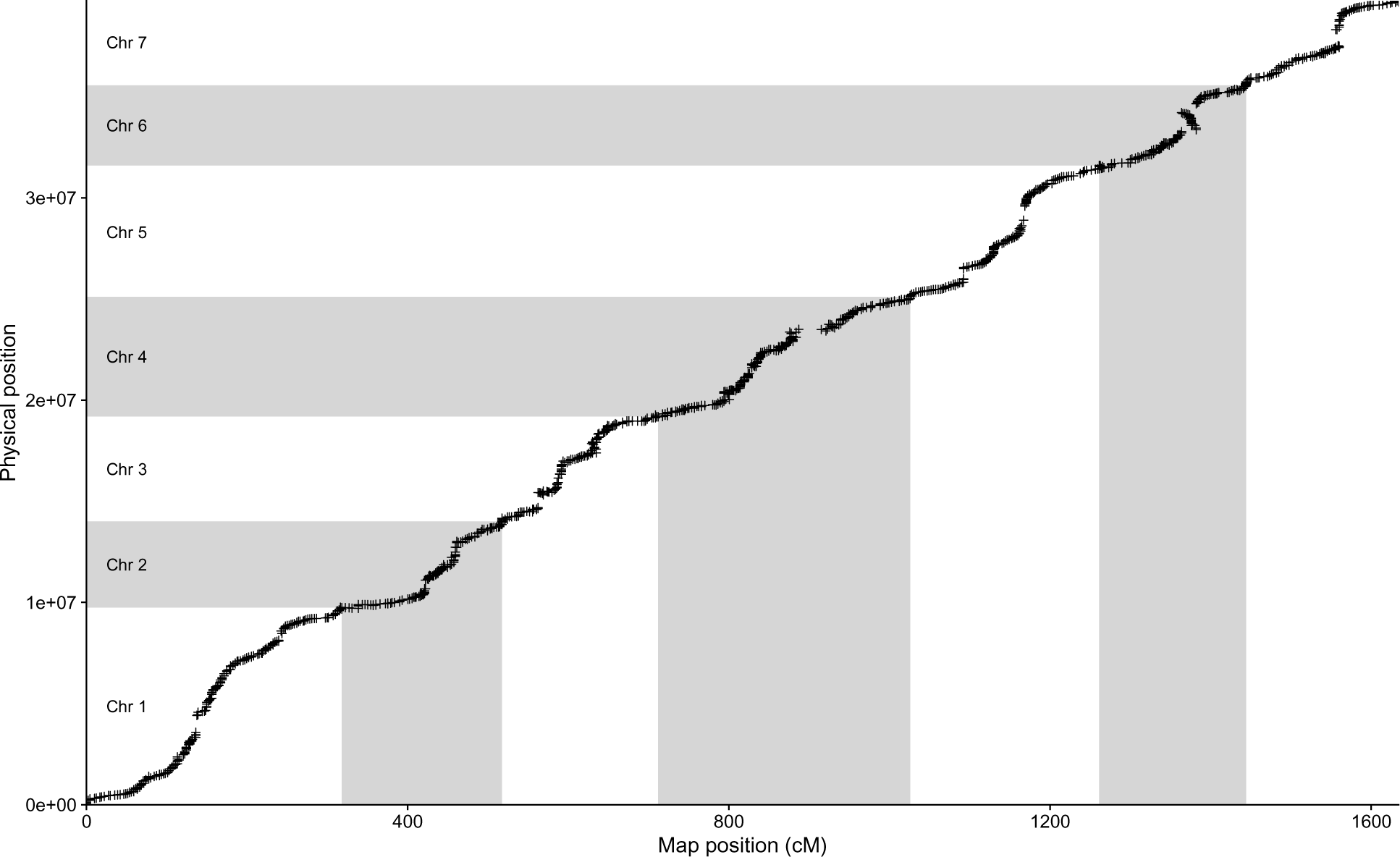
An example of comparing marker order on the genetic map to the physical reference genome for family A. There are only a few cases where markers may be in the wrong order. On chromosome 6 there appears to be an inversion relative to the reference genome.

**Figure S4:**
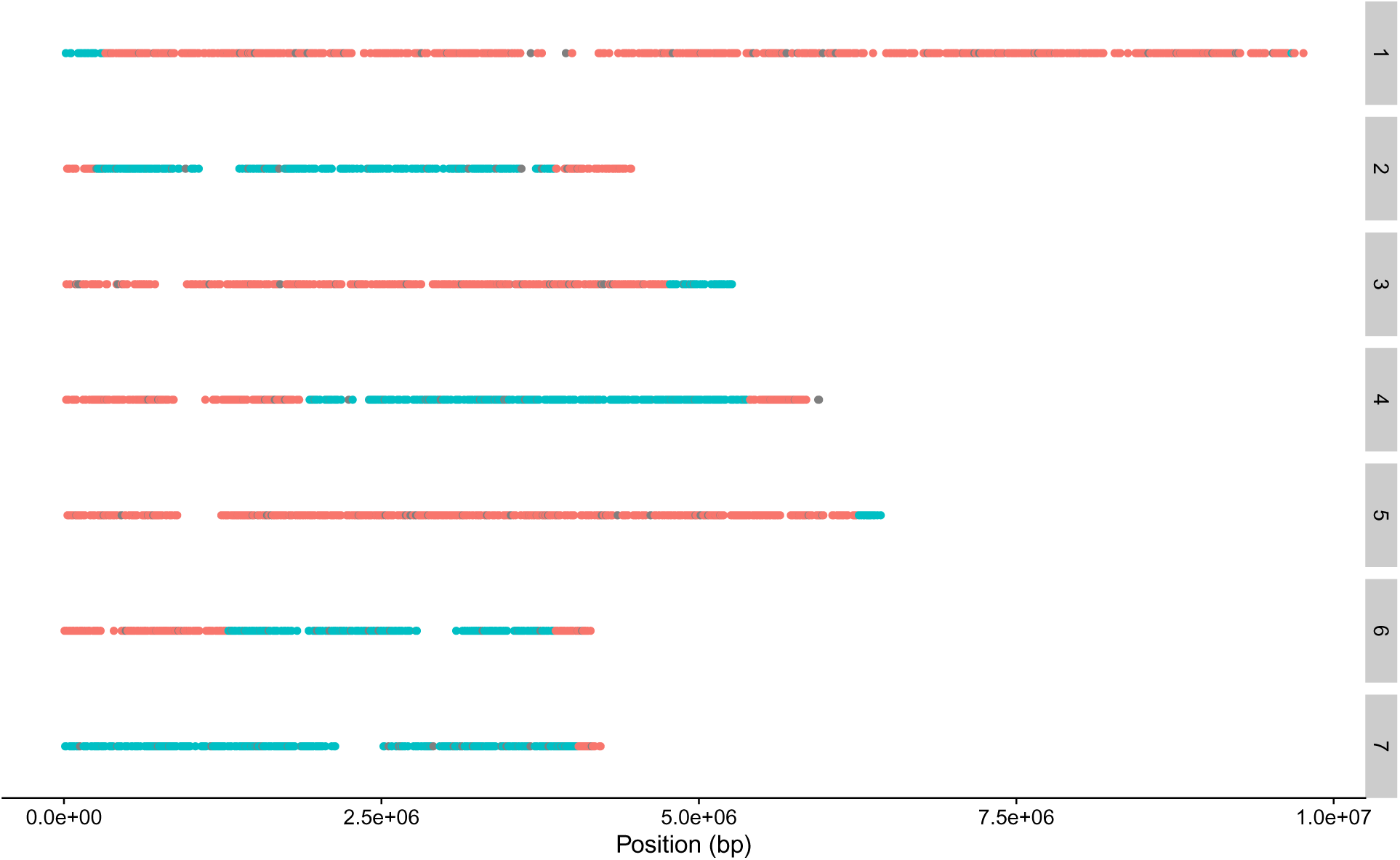
An example of strain A2 with segments of the genome inherited from the two parents: 10886 (red) and 10948 (blue). Grey points are markers with missing data. Facet labeling shows chromosome numbers.

**Figure S5:**
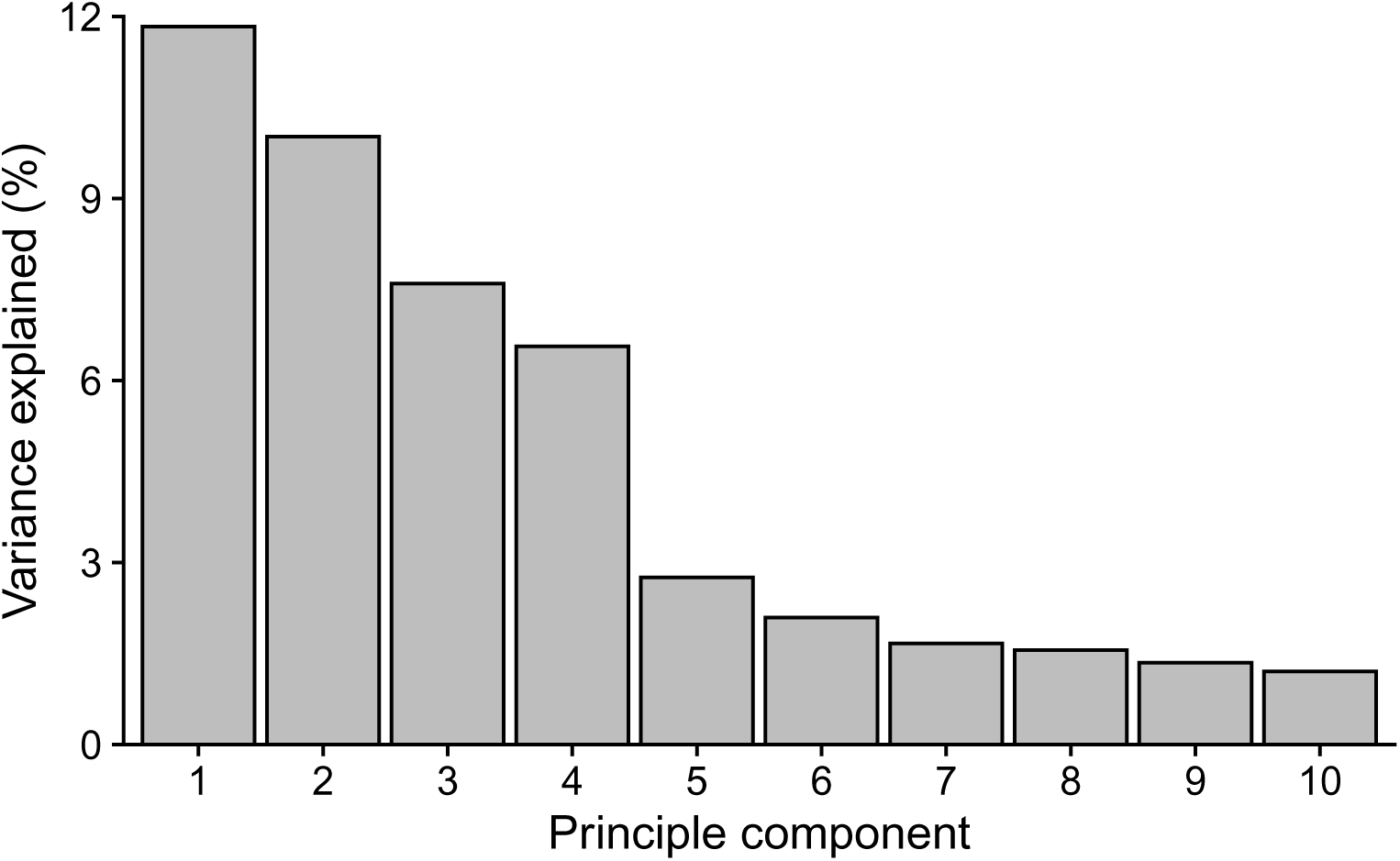
The amount of variance explained by the different components when principle component analysis was performed for the SNP data of the nested association mapping population.

**Figure S6:**
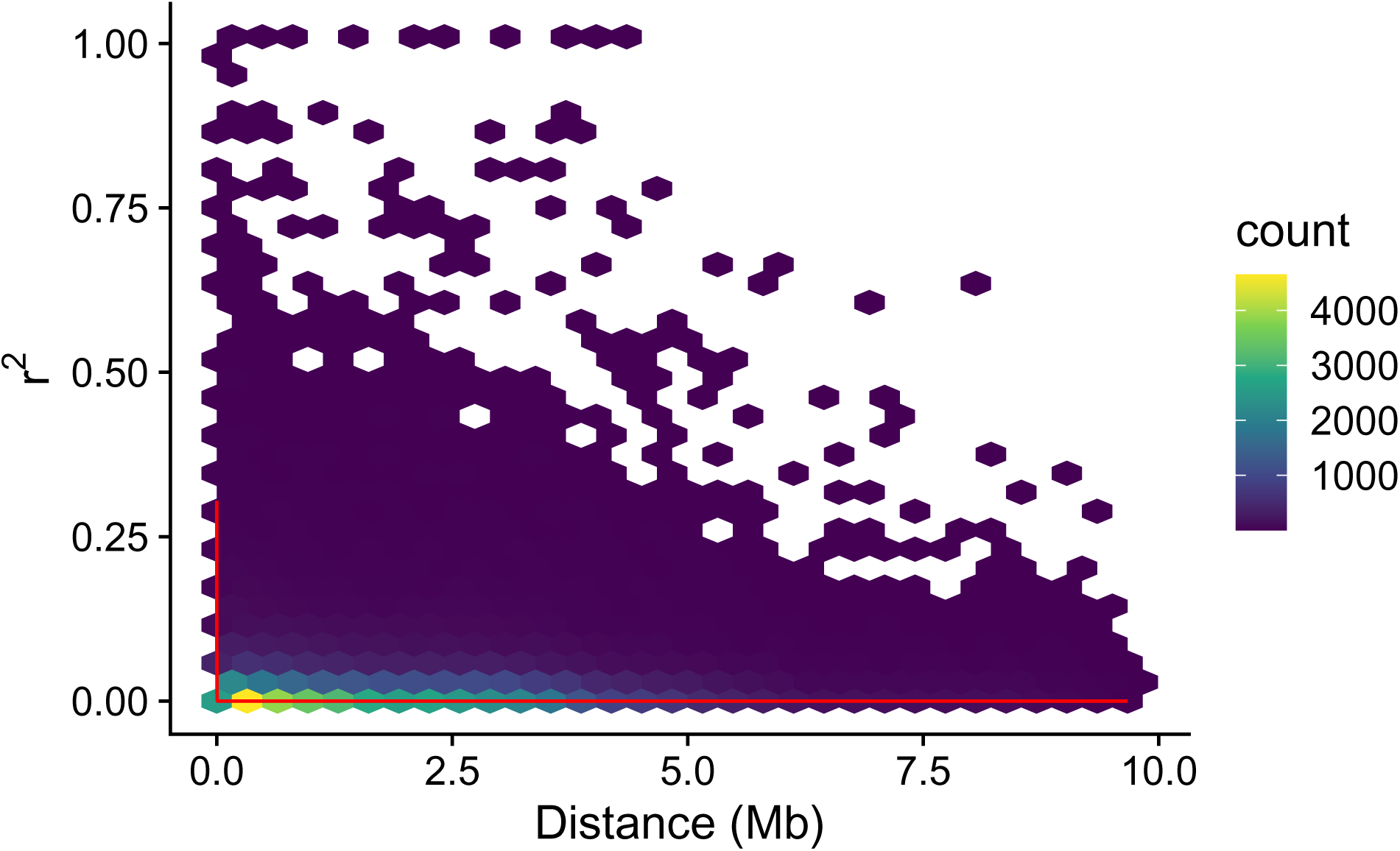
Decay of linkage disequilibrium over long distances across the seven chromosomes of *N. crassa*. We sampled 10 000 pairwise SNP combinations at random from each chromosome, *n* = 70 000 in total. For plotting data has been binned into hexes. Red line shows exponential decay fit.

**Figure S7:**
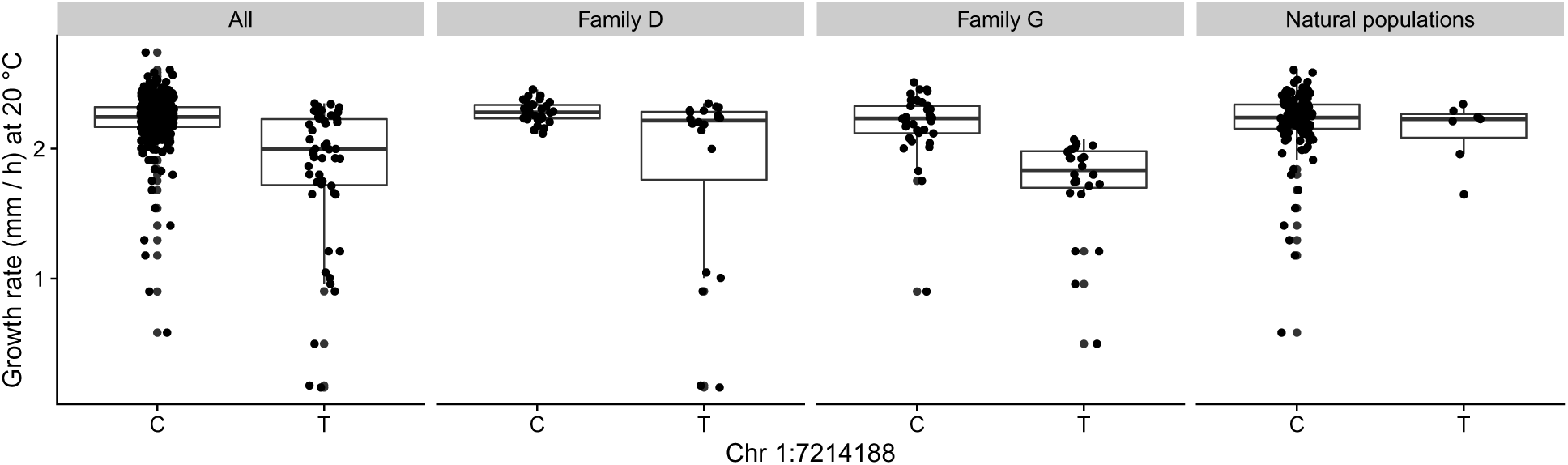
Segregation of SNPs significant at 20 °C.

**Figure S8:**
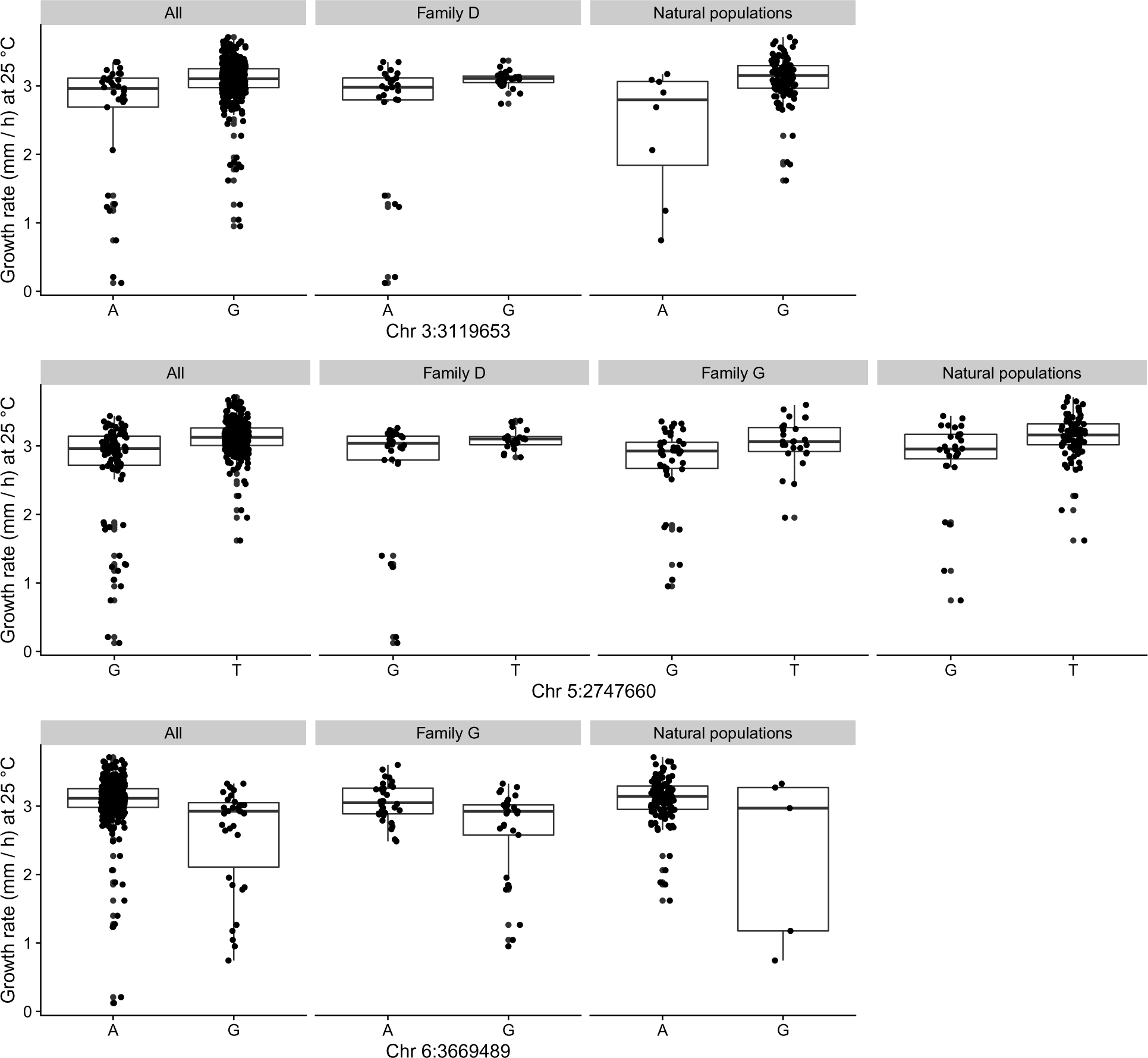
Segregation of SNPs significant at 25 °C.

**Figure S9:**
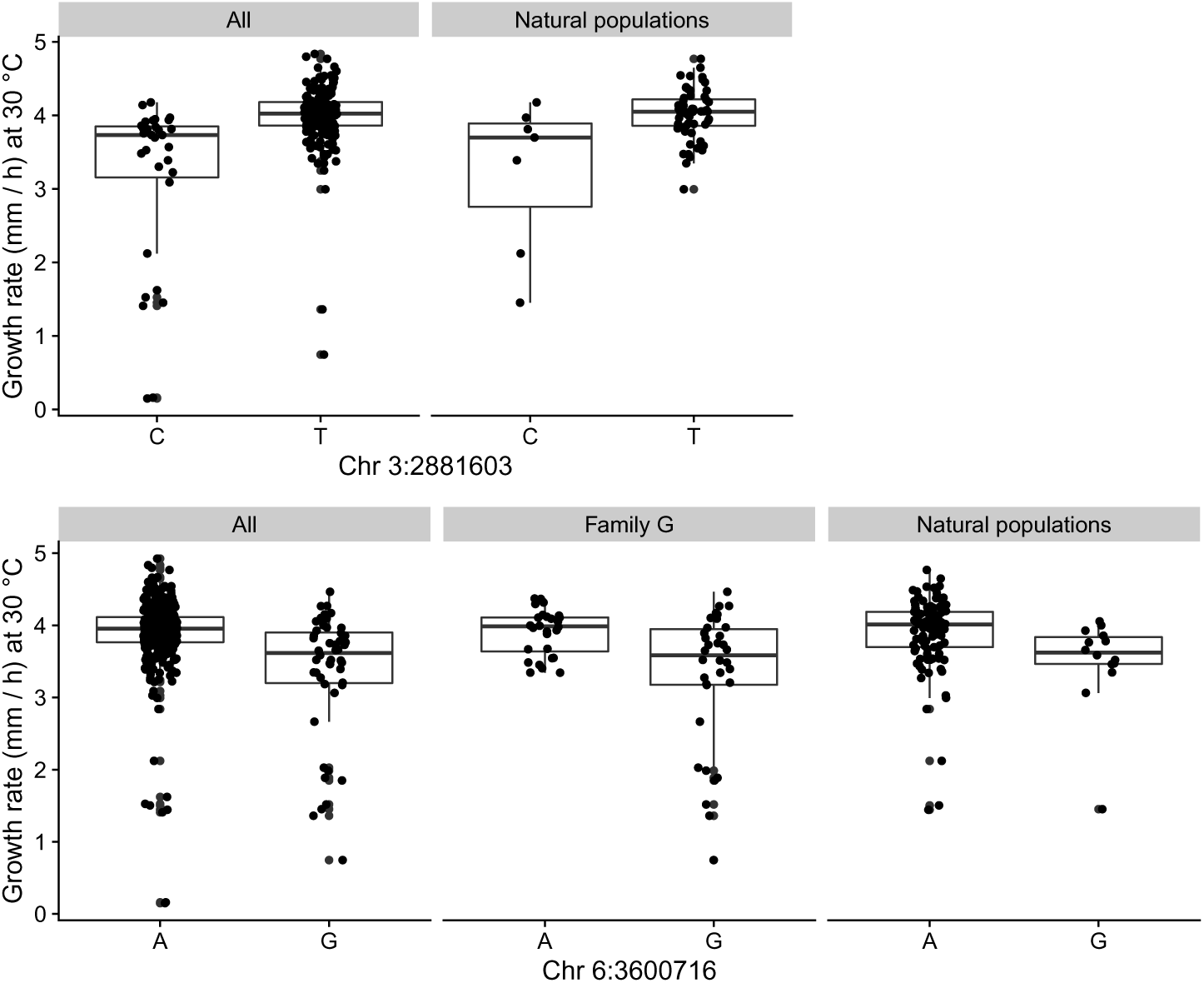
Segregation of SNPs significant at 30 °C.

**Figure S10:**
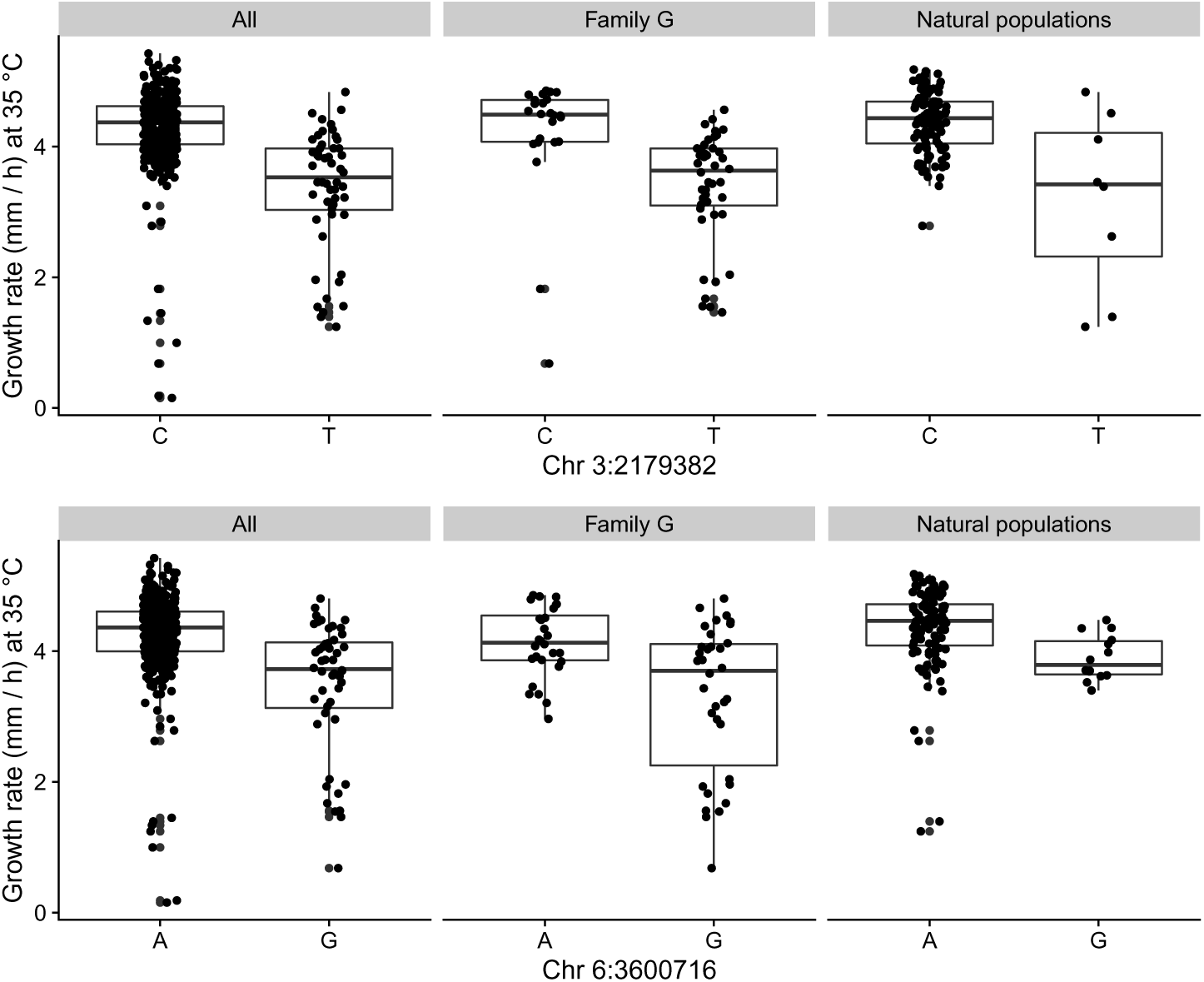
Segregation of SNPs significant at 35 °C.

**Figure S11:**
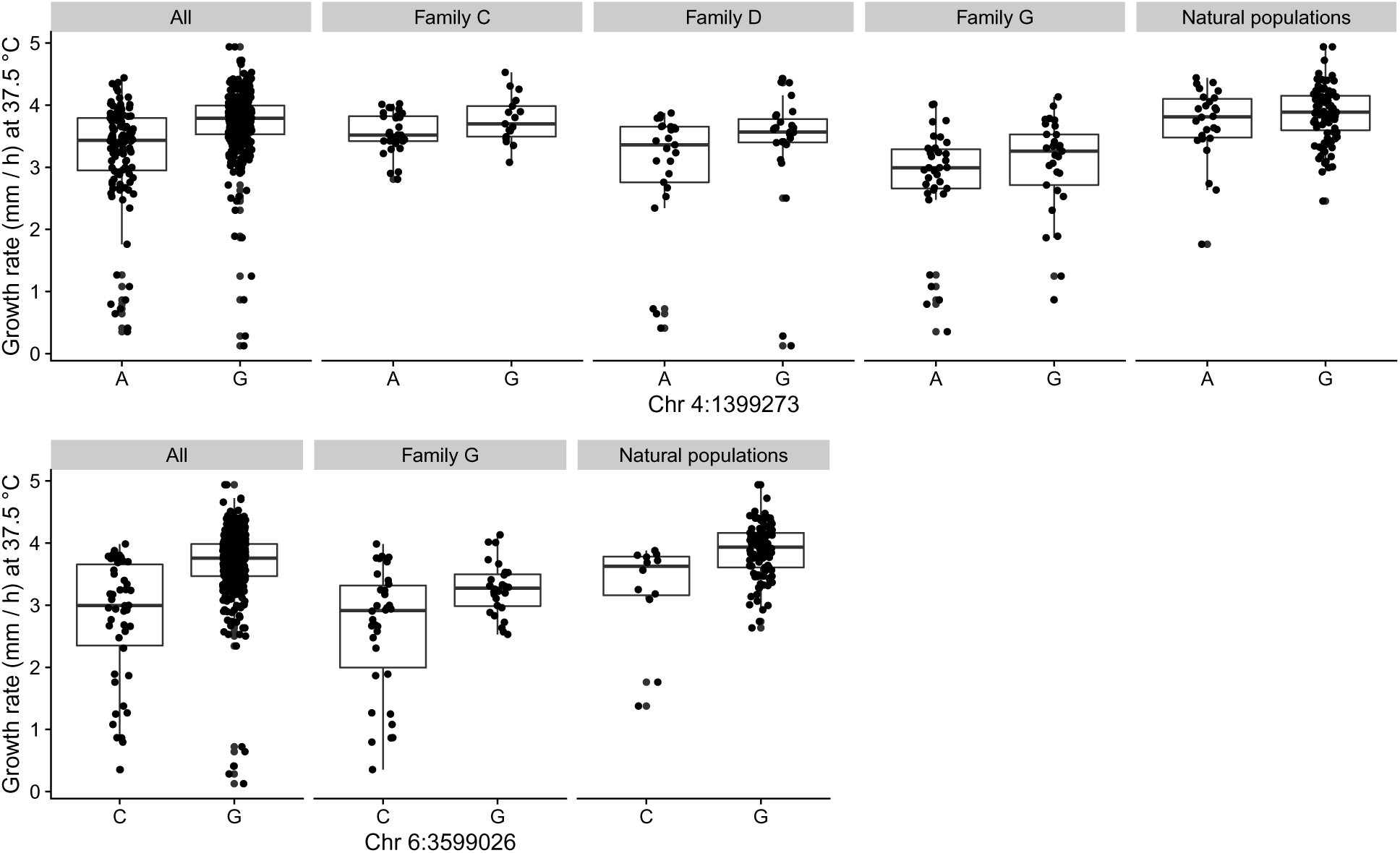
Segregation of SNPs significant at 37.5 °C.

**Figure S12:**
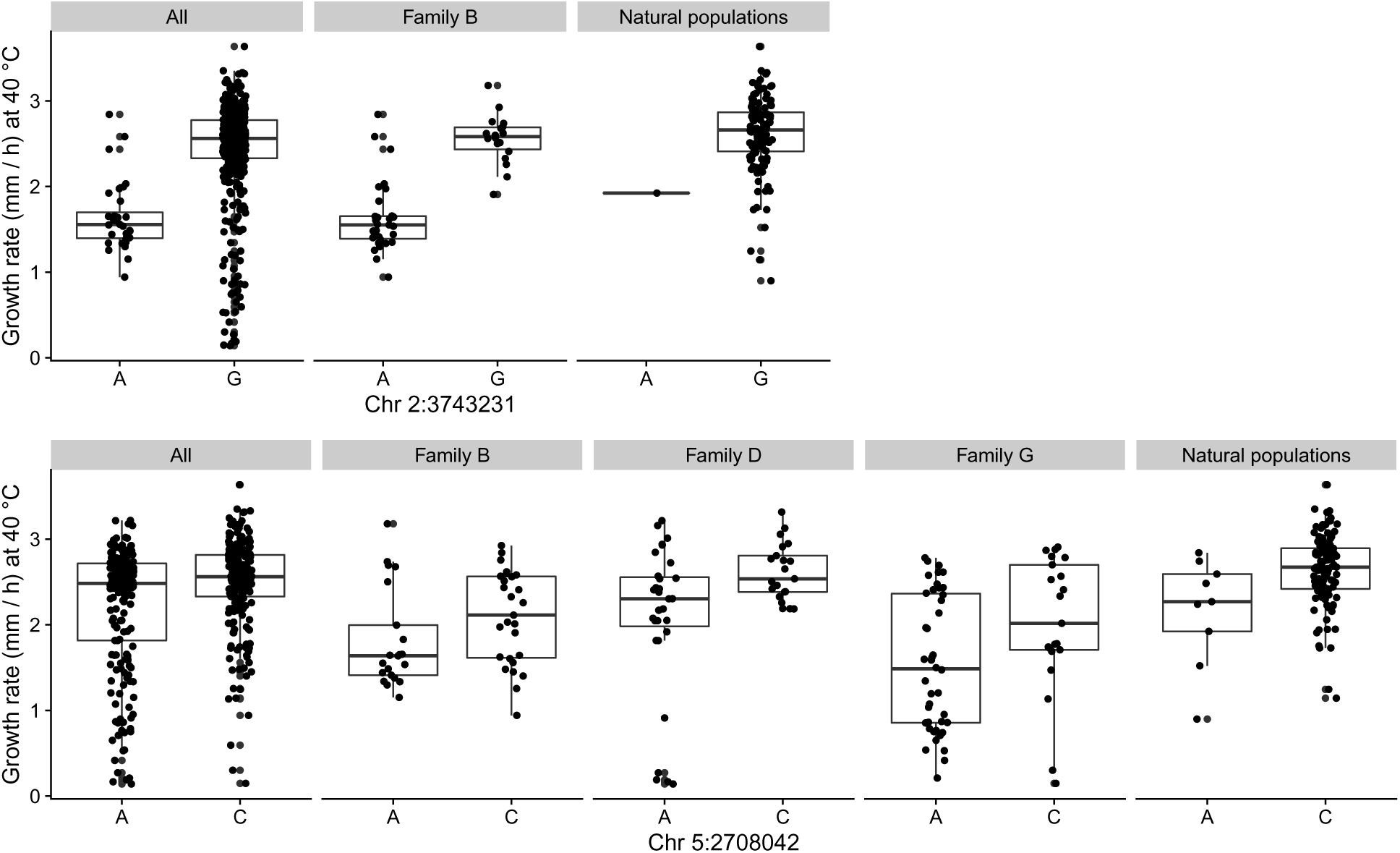
Segregation of SNPs significant at 40 °C.

**Figure S13:**
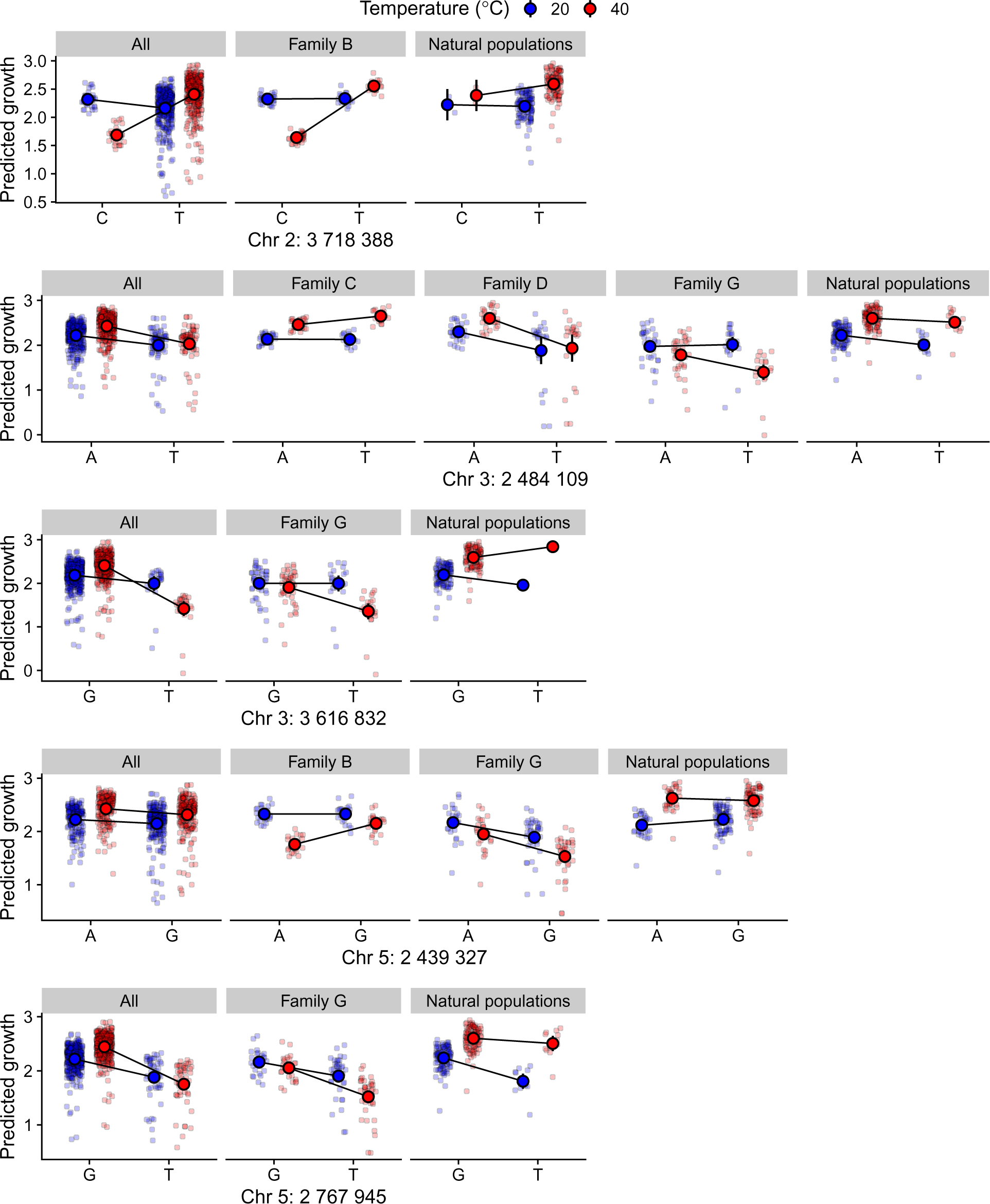
Allelic effects for SNPs that were associated with TPC breadth and had no evidence for trade-offs between 20 and 40 °C.

### Supplementary Tables

**Table S1:**
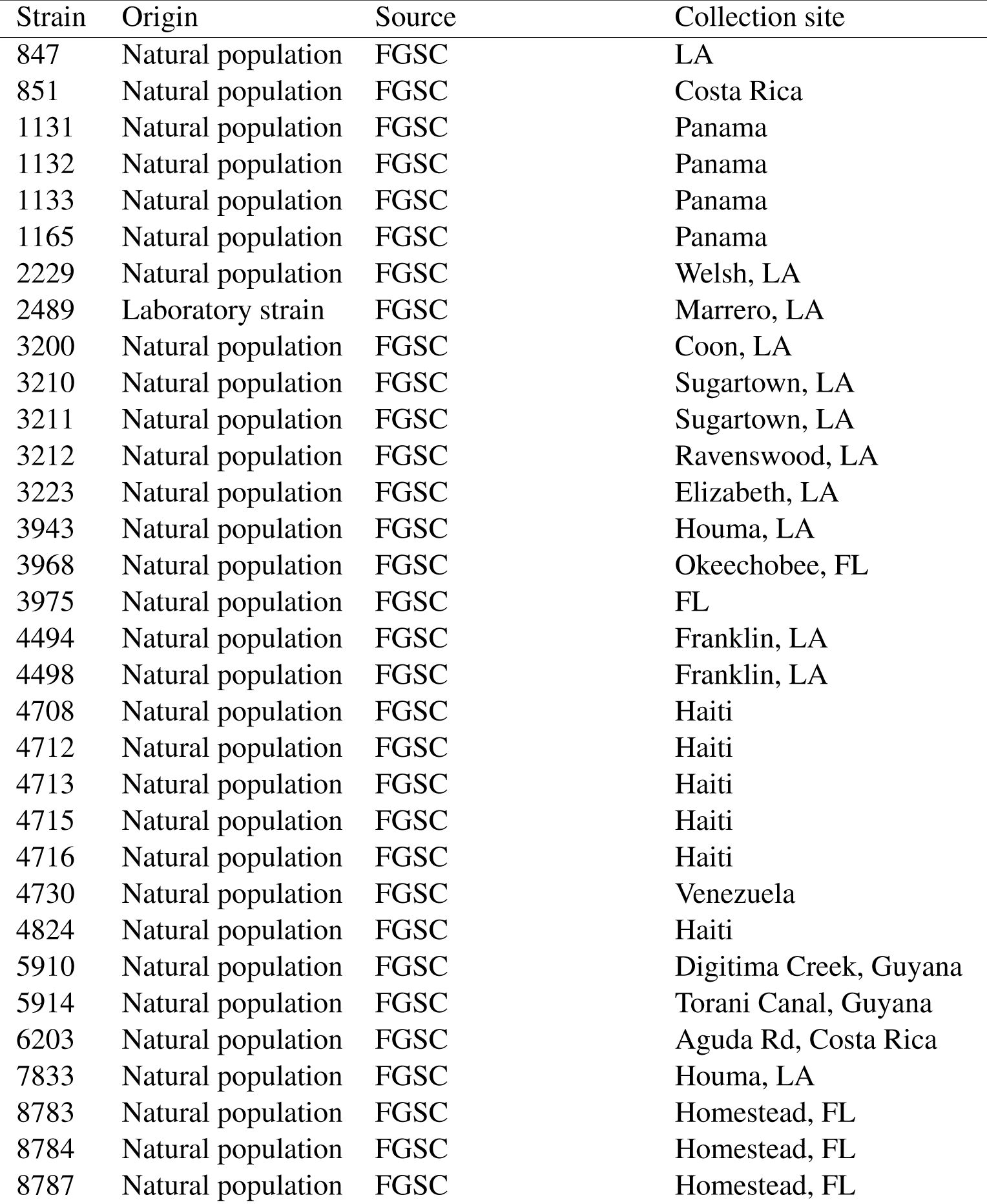
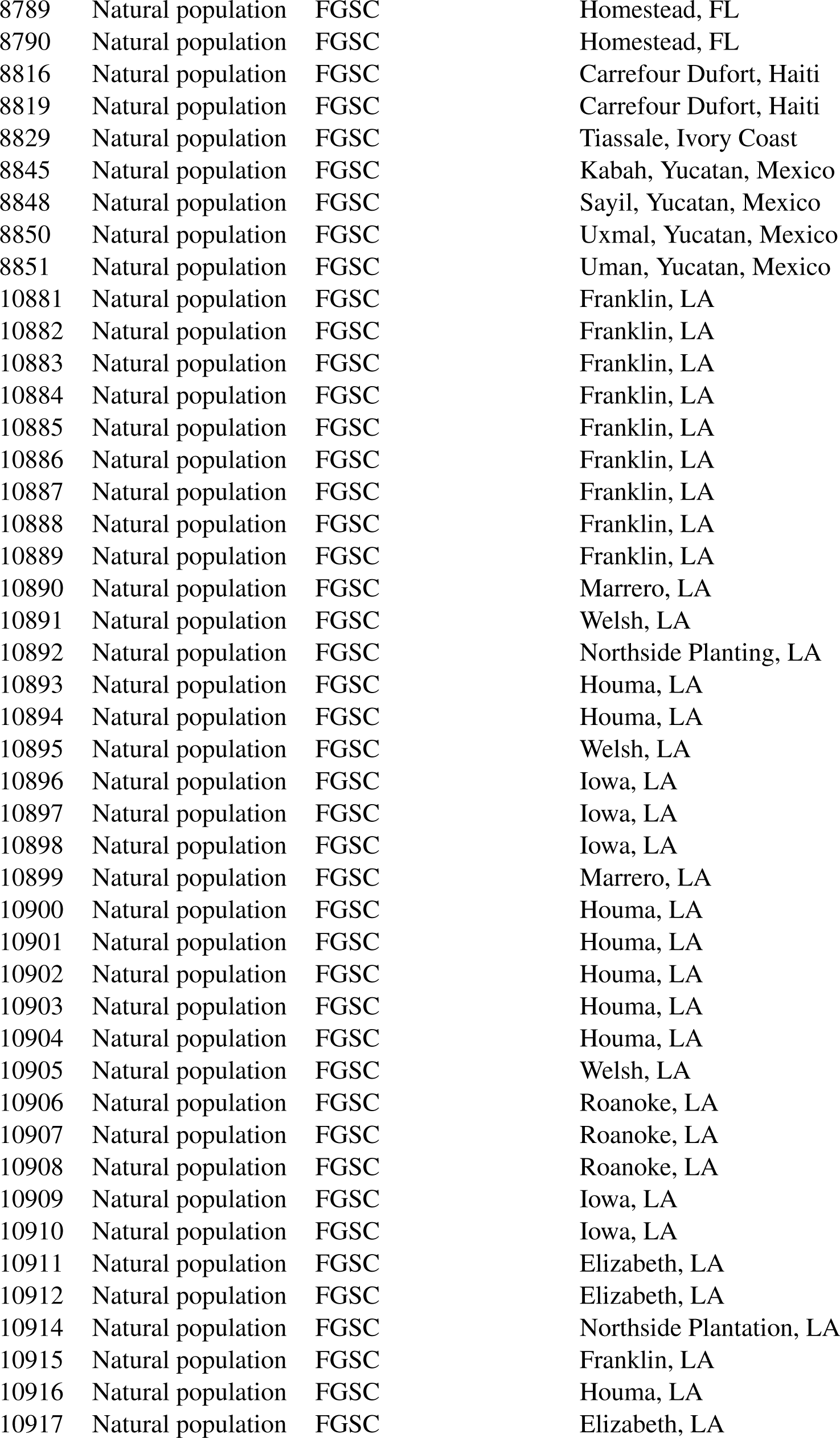
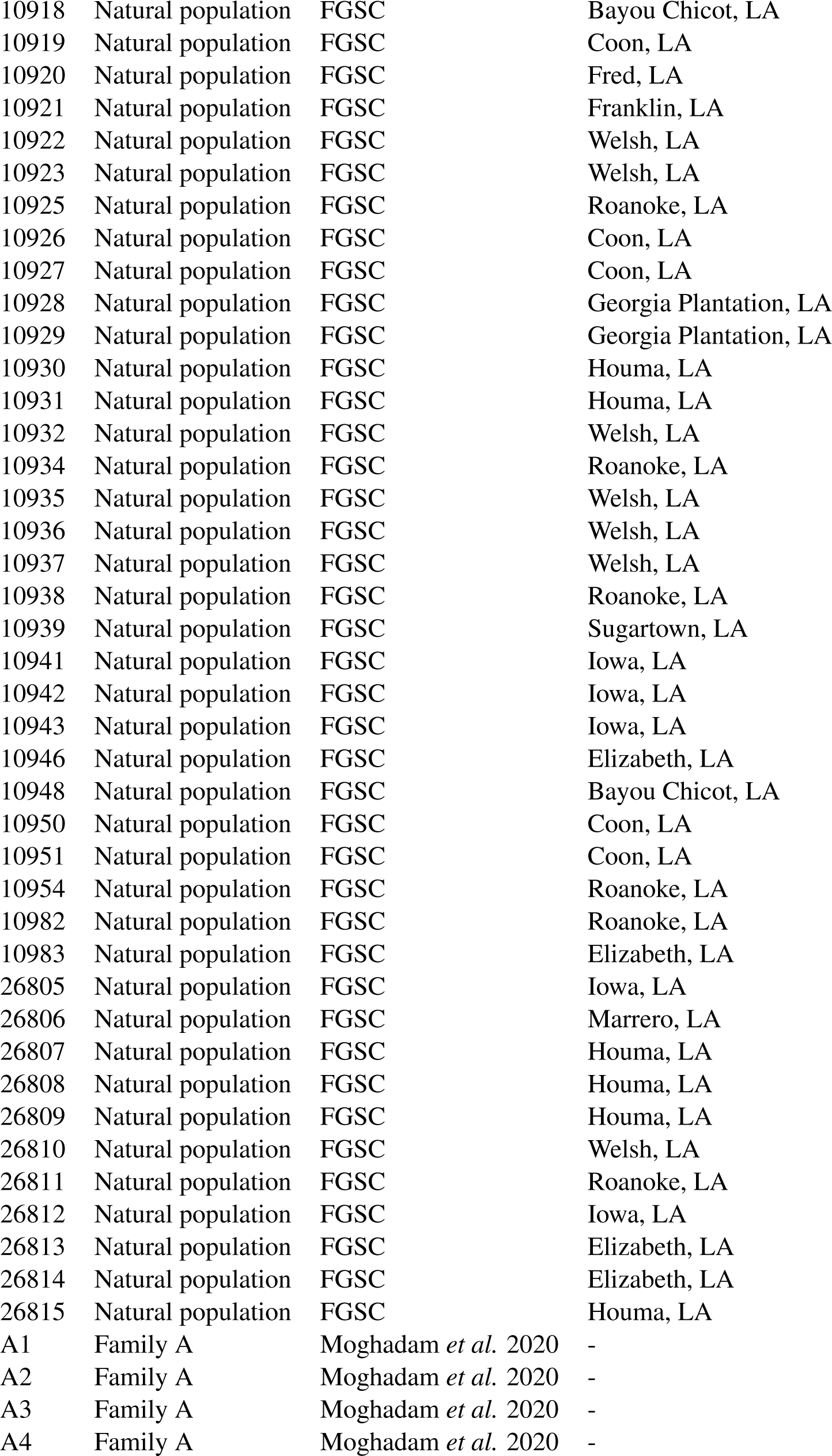
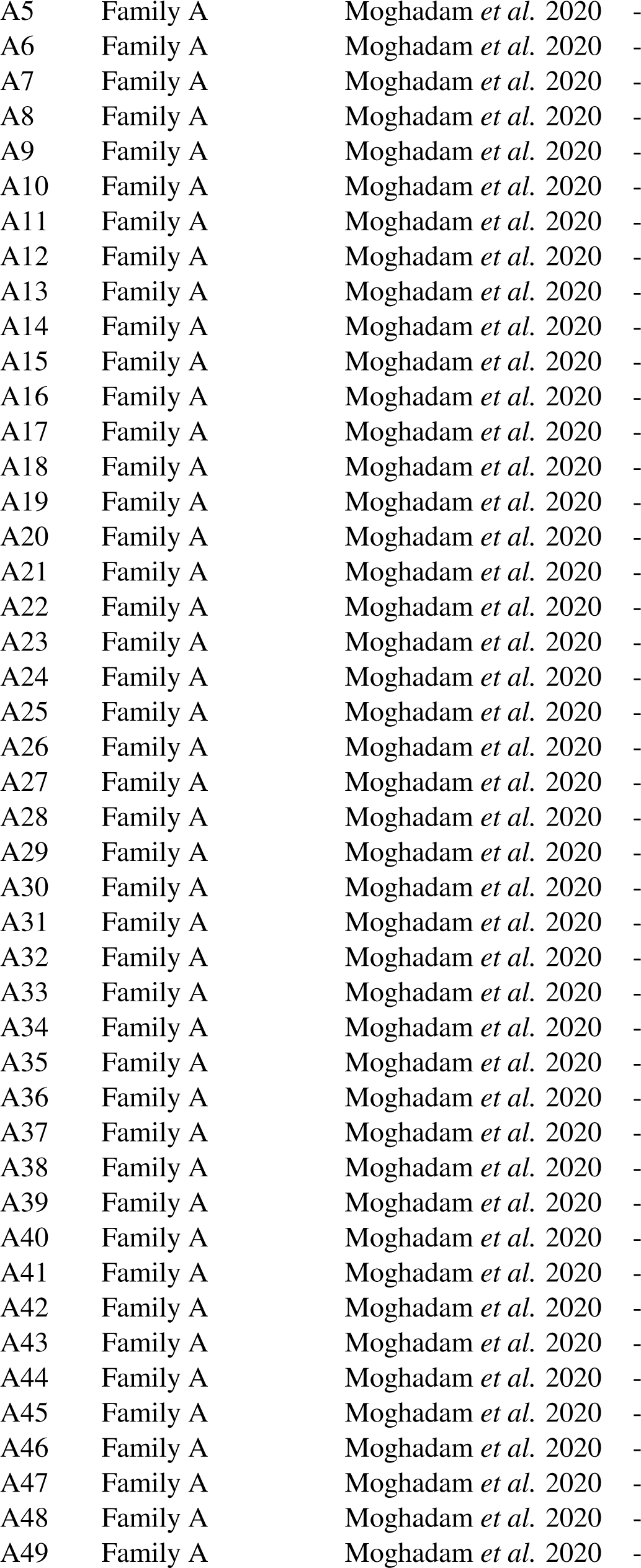
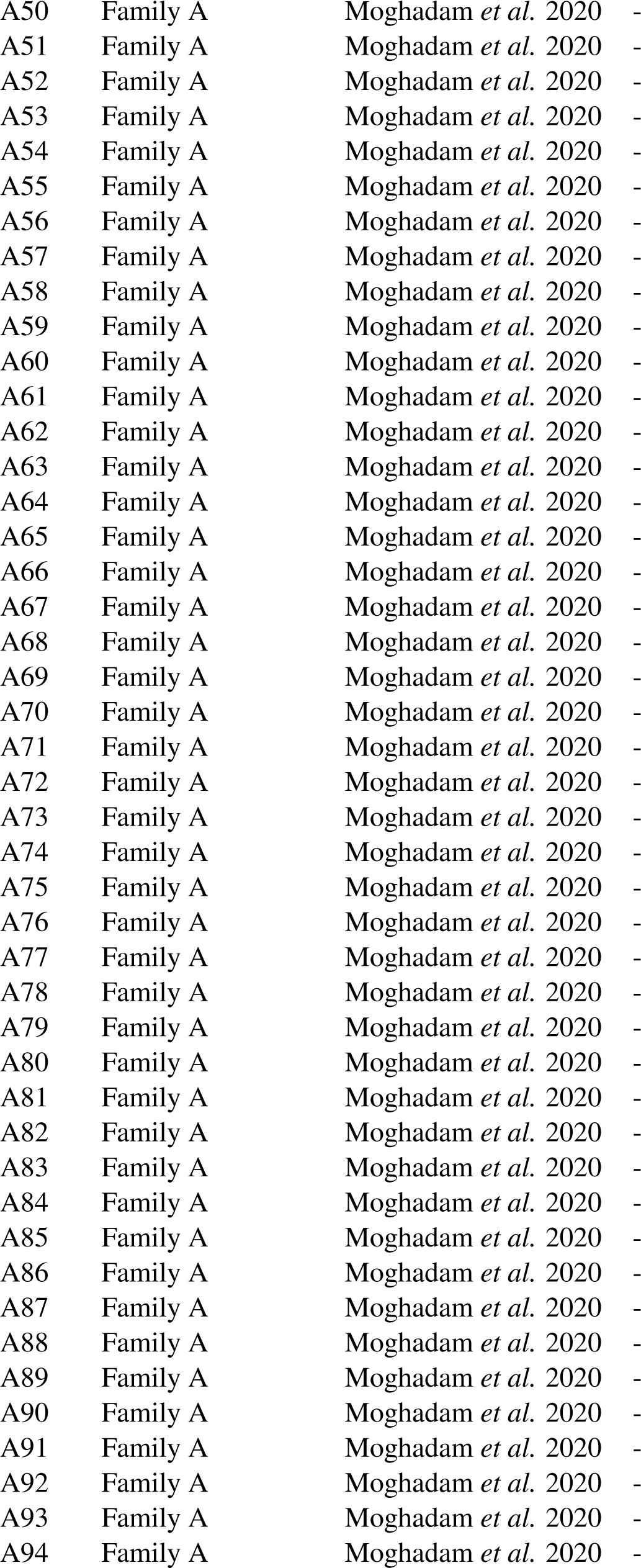
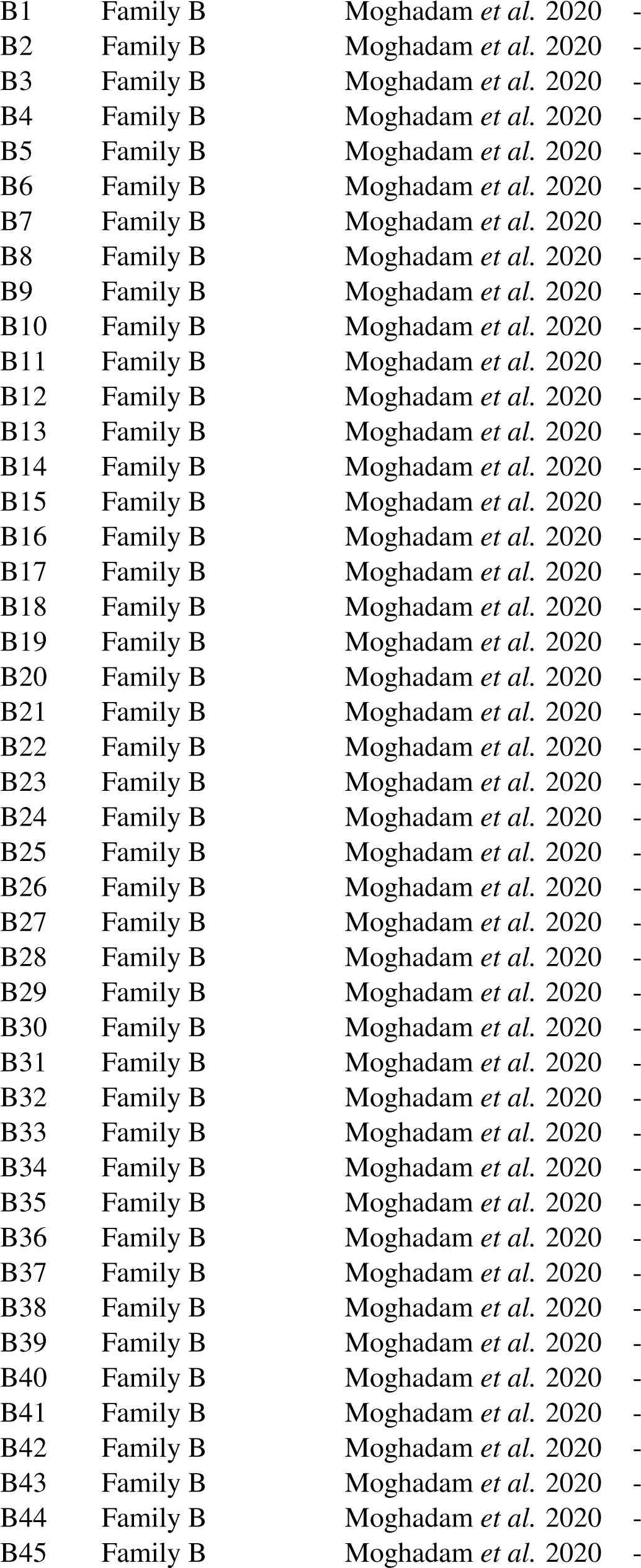
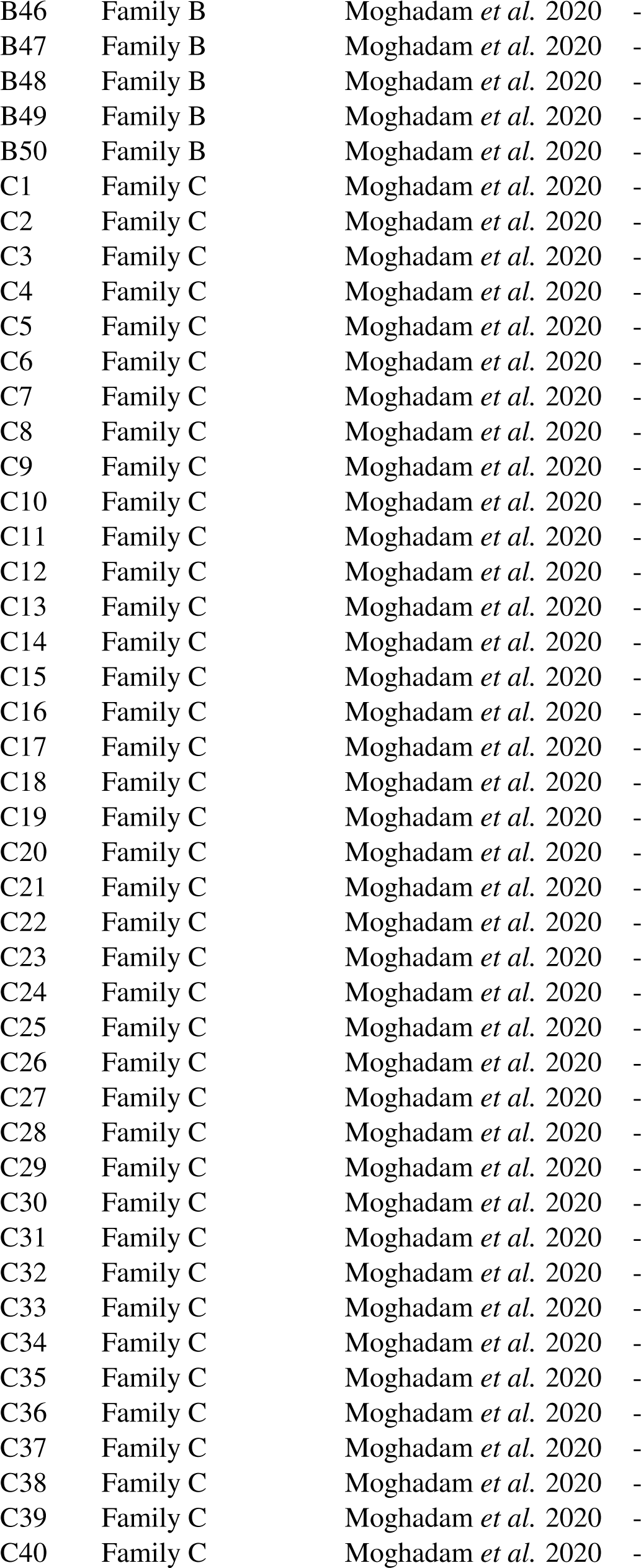
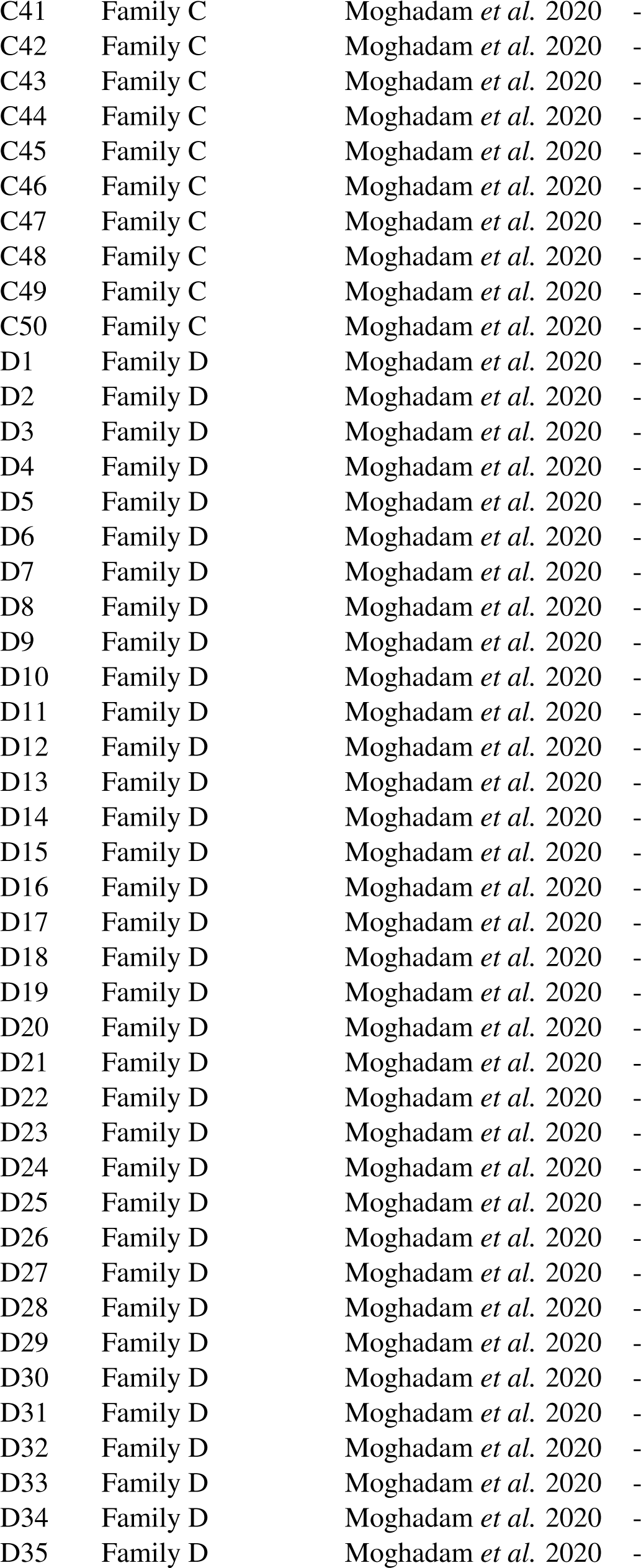
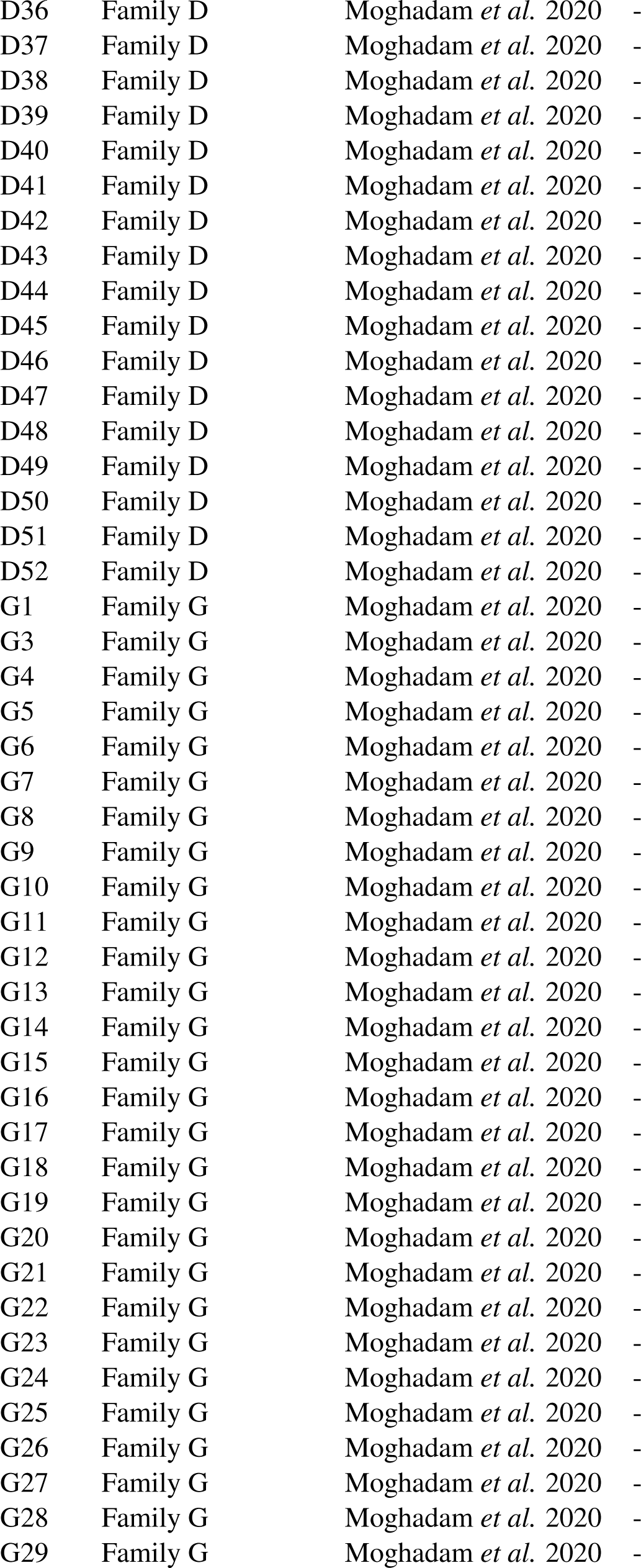
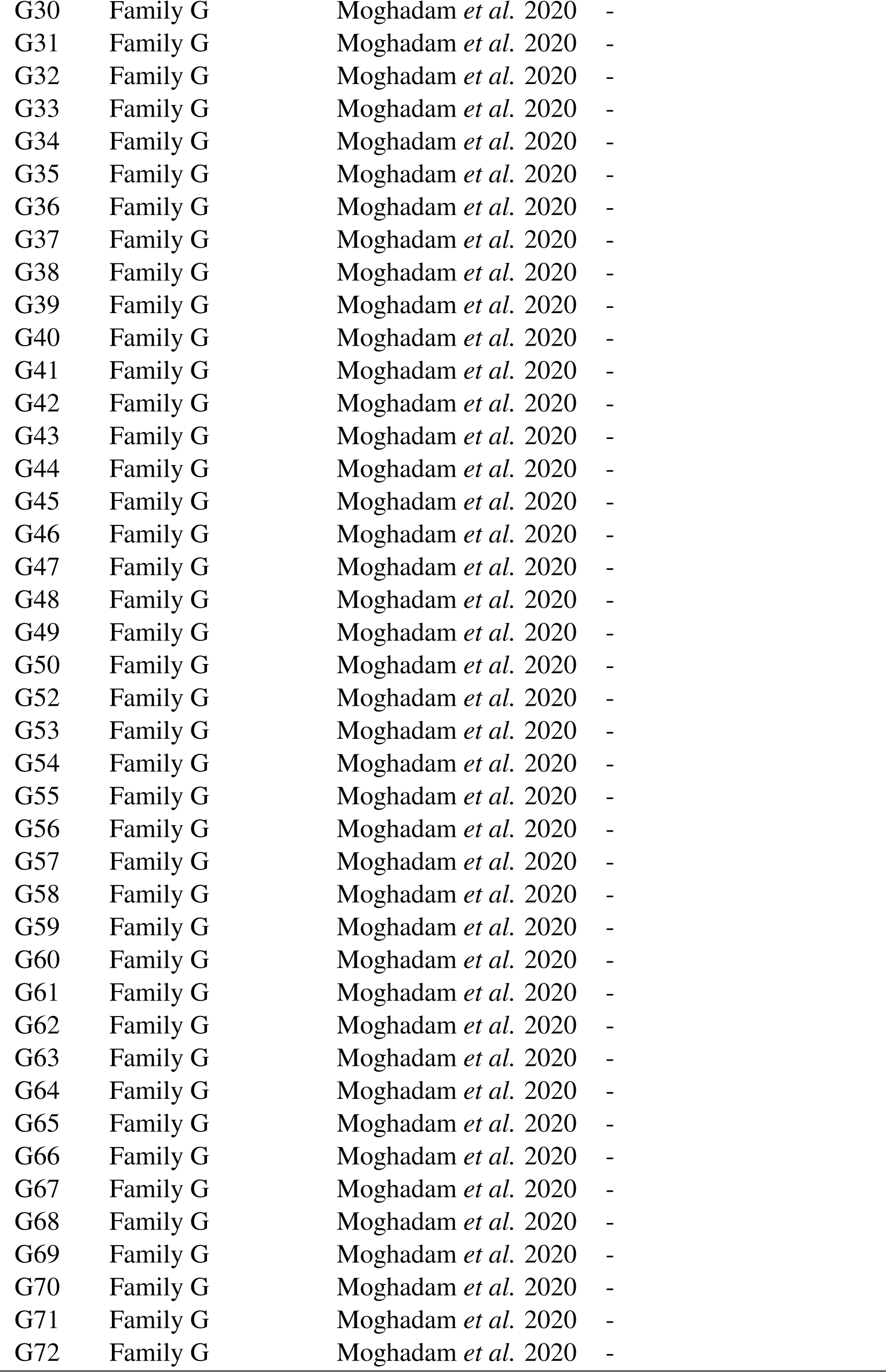
List of strains. Column origin indicates whether the strain was sampled from a natural population or if it was from a family obtained by crossing two natural strains. Column source indicates which strains were obtained from the Fungal Genetics Stock Center (FGSC) and which were generated in Moghadam *et al*. 2020. LA = Louisiana, USA. FL = Florida, USA. Strains 10948 and 10886 are parents of family A, 10932 and 1165 are parents of family B, 4498 and 8816 are parents of family C, 3223 and 8845 are parents of family D, and 10904 and 851 are parents of family G.

**Table S2:** Summary of alignment metrics for re-sequenced strains. Strain is the FGSC strain number, synonym is the alternative strain name, mapped reads show the number of reads mapped to the reference genome, coverage shows how much of the reference genome was covered, depth is the weighted average of sequencing depth, and source indicates whether the strain was sequenced in this study or if the reads were obtained from previous studies.

| Strain | Synonym | Mapped reads | Coverage (%) | Depth | Source |
| --- | --- | --- | --- | --- | --- |
| 847 |  | 15425851 | 97 | 55 | This study |
| 851 |  | 13209563 | 92 | 45 | Villalba de la Peña *et al.* (2023) |
| 1131 |  | 17337934 | 93 | 58 | Villalba de la Peña *et al.* (2023) |
| 1132 |  | 16674240 | 93 | 57 | This study |
| 1133 |  | 14784777 | 93 | 50 | Villalba de la Peña *et al.* (2023) |
| 1165 |  | 15409953 | 93 | 52 | Villalba de la Peña *et al.* (2023) |
| 2229 |  | 15324074 | 94 | 53 | This study |
| 3200 |  | 15151373 | 94 | 52 | This study |
| 3210 |  | 13946131 | 94 | 48 | Villalba de la Peña *et al.* (2023) |
| 3211 |  | 14185248 | 93 | 48 | Villalba de la Peña *et al.* (2023) |
| 3212 |  | 17827161 | 94 | 61 | This study |
| 3223 |  | 14358887 | 94 | 49 | Villalba de la Peña *et al.* (2023) |
| 3943 |  | 143996355 | 97 | 170 | Zhao *et al.* (2015) |
| 3968 |  | 16916617 | 93 | 58 | This study |
| 3975 |  | 13947120 | 92 | 45 | Villalba de la Peña *et al.* (2023) |
| 4494 |  | 19630480 | 100 | 71 | This study |
| 4498 |  | 16351032 | 97 | 58 | This study |
| 4708 |  | 14636965 | 95 | 49 | Villalba de la Peña *et al.* (2023) |
| 4712 |  | 12929356 | 93 | 43 | Villalba de la Peña *et al.* (2023) |
| 4713 |  | 20045479 | 93 | 69 | This study |
| 4715 |  | 19120259 | 93 | 65 | This study |
| 4716 |  | 18759224 | 93 | 63 | Villalba de la Peña *et al.* (2023) |
| 4730 |  | 15865760 | 93 | 53 | Villalba de la Peña *et al.* (2023) |
| 4824 |  | 13207415 | 93 | 44 | Villalba de la Peña *et al.* (2023) |
| 5910 |  | 12714008 | 93 | 42 | Villalba de la Peña *et al.* (2023) |
| 5914 |  | 14781732 | 92 | 50 | This study |
| 6203 |  | 13538150 | 92 | 46 | Villalba de la Peña *et al.* (2023) |
| 7833 | P4448 | 19807530 | 94 | 68 | This study |
| 8783 |  | 15743318 | 93 | 52 | Villalba de la Peña *et al.* (2023) |
| 8784 |  | 17605429 | 94 | 60 | This study |
| 8787 |  | 17652705 | 93 | 60 | This study |
| 8789 |  | 17625386 | 93 | 60 | This study |
| 8790 |  | 13456207 | 92 | 46 | Villalba de la Peña *et al.* (2023) |
| 8816 |  | 15765804 | 93 | 54 | Villalba de la Peña *et al.* (2023) |
| 8819 |  | 14047916 | 92 | 47 | Villalba de la Peña *et al.* (2023) |
| 8829 |  | 17445062 | 92 | 59 | This study |
| 8845 |  | 16226721 | 93 | 55 | Villalba de la Peña *et al.* (2023) |
| 8848 |  | 14429838 | 93 | 49 | This study |

Table S2 – Continued
| Strain | Synonym | Mapped reads | Coverage (%) | Depth | Source |
| --- | --- | --- | --- | --- | --- |
| 8850 |  | 12959487 | 92 | 44 | Villalba de la Peña *et al.* (2023) |
| 8851 |  | 17393784 | 93 | 59 | This study |
| 10881 |  | 16941381 | 94 | 58 | This study |
| 10882 |  | 30800959 | 95 | 71 | Zhao *et al.* (2015) |
| 10883 |  | 17141000 | 94 | 59 | Zhao *et al.* (2015) |
| 10884 |  | 18031050 | 95 | 62 | Zhao *et al.* (2015) |
| 10885 |  | 15768753 | 94 | 54 | This study |
| 10886 |  | 15527529 | 93 | 53 | Villalba de la Peña *et al.* (2023) |
| 10887 |  | 15501471 | 94 | 53 | This study |
| 10888 |  | 17851313 | 94 | 61 | This study |
| 10889 |  | 19218000 | 95 | 66 | This study |
| 10890 |  | 17491081 | 100 | 63 | This study |
| 10891 |  | 16977210 | 100 | 61 | This study |
| 10892 |  | 32427959 | 95 | 71 | Zhao *et al.* (2015) |
| 10893 |  | 18173287 | 94 | 63 | This study |
| 10894 |  | 16601337 | 94 | 56 | This study |
| 10895 |  | 17816777 | 100 | 64 | This study |
| 10896 |  | 16169786 | 100 | 58 | This study |
| 10897 |  | 17123714 | 93 | 58 | This study |
| 10898 |  | 15849395 | 93 | 54 | This study |
| 10899 |  | 18243325 | 94 | 62 | This study |
| 10900 |  | 16972252 | 84 | 54 | This study |
| 10901 |  | 19722355 | 93 | 67 | This study |
| 10902 |  | 23980217 | 95 | 82 | This study |
| 10903 |  | 19114408 | 94 | 66 | This study |
| 10904 |  | 14997206 | 95 | 51 | Villalba de la Peña *et al.* (2023) |
| 10905 |  | 18917865 | 94 | 64 | This study |
| 10906 |  | 15257473 | 94 | 52 | Villalba de la Peña *et al.* (2023) |
| 10907 |  | 32506031 | 95 | 71 | Zhao *et al.* (2015) |
| 10908 |  | 14173072 | 94 | 49 | Villalba de la Peña *et al.* (2023) |
| 10909 |  | 17240390 | 94 | 59 | This study |
| 10910 |  | 17022398 | 94 | 58 | This study |
| 10911 |  | 18577891 | 94 | 64 | This study |
| 10912 |  | 15107747 | 94 | 52 | Villalba de la Peña *et al.* (2023) |
| 10914 |  | 30425792 | 95 | 67 | Zhao *et al.* (2015) |
| 10915 |  | 24565381 | 96 | 55 | Zhao *et al.* (2015) |
| 10916 |  | 24695915 | 94 | 85 | This study |
| 10917 |  | 11183264 | 94 | 38 | This study |
| 10918 |  | 40504581 | 95 | 89 | Zhao *et al.* (2015) |
| 10919 |  | 18812053 | 94 | 64 | This study |
| 10920 |  | 15056084 | 94 | 52 | This study |
| 10921 |  | 17963443 | 95 | 62 | This study |
| 10922 |  | 19083647 | 93 | 65 | This study |
| 10923 |  | 16519837 | 94 | 57 | Villalba de la Peña *et al.* (2023) |
| 10925 |  | 37150140 | 95 | 86 | Zhao *et al.* (2015) |

Table S2 – Continued
| Strain | Synonym | Mapped reads | Coverage (%) | Depth | Source |
| --- | --- | --- | --- | --- | --- |
| 10926 |  | 32962766 | 95 | 73 | Zhao *et al.* (2015) |
| 10927 |  | 17862238 | 95 | 40 | Zhao *et al.* (2015) |
| 10928 |  | 15315502 | 94 | 52 | Villalba de la Peña *et al.* (2023) |
| 10929 |  | 18766216 | 94 | 64 | This study |
| 10930 |  | 16790516 | 94 | 58 | This study |
| 10931 |  | 16445241 | 94 | 56 | This study |
| 10932 |  | 13675057 | 94 | 47 | Villalba de la Peña *et al.* (2023) |
| 10934 |  | 17718497 | 94 | 60 | This study |
| 10935 |  | 22403064 | 95 | 51 | Zhao *et al.* (2015) |
| 10936 |  | 18996835 | 94 | 65 | This study |
| 10937 |  | 29871313 | 97 | 69 | Zhao *et al.* (2015) |
| 10938 |  | 18613612 | 94 | 64 | This study |
| 10939 |  | 17427259 | 95 | 59 | This study |
| 10941 |  | 15700310 | 94 | 54 | This study |
| 10942 |  | 17227423 | 96 | 59 | This study |
| 10943 |  | 33098168 | 95 | 73 | Zhao *et al.* (2015) |
| 10946 |  | 17632666 | 93 | 59 | Villalba de la Peña *et al.* (2023) |
| 10948 |  | 14115769 | 94 | 48 | Villalba de la Peña *et al.* (2023) |
| 10950 |  | 14490901 | 100 | 52 | Villalba de la Peña *et al.* (2023) |
| 10951 |  | 13650273 | 94 | 46 | Villalba de la Peña *et al.* (2023) |
| 10954 |  | 25167395 | 100 | 91 | This study |
| 10982 |  | 16317316 | 96 | 56 | This study |
| 10983 |  | 38636650 | 95 | 87 | Zhao *et al.* (2015) |
| 26805 | P4451 | 17359255 | 94 | 60 | This study |
| 26806 | P4452 | 172181523 | 95 | 206 | Zhao *et al.* (2015) |
| 26807 | P4457 | 17847184 | 94 | 60 | This study |
| 26808 | P4459 | 15529408 | 94 | 53 | This study |
| 26809 | P4463 | 21876017 | 95 | 49 | Zhao *et al.* (2015) |
| 26810 | P4468 | 21305398 | 96 | 47 | Zhao *et al.* (2015) |
| 26811 | P4471 | 25006742 | 96 | 56 | Zhao *et al.* (2015) |
| 26812 | P4472 | 17162696 | 94 | 59 | This study |
| 26813 | P4479 | 22395100 | 95 | 49 | Zhao *et al.* (2015) |
| 26814 | P4486 | 21584057 | 94 | 74 | This study |
| 26815 | P4496 | 19603506 | 94 | 68 | This study |

